# Functional immune profiling reveals CD4^+^ T cell dysregulation associated with coeliac disease

**DOI:** 10.1101/2025.06.06.658209

**Authors:** Anthony J. Farchione, HoChan Cheon, David Vremec, Julika Neumann, Gwenny M. Verstappen, Melinda Y. Hardy, Mai B. Margetts, Lauren J. Howson, Lee M. Henneken, Maureen Forde, Jason A. Tye-Din, Susanne Heinzel, Philip D. Hodgkin, Vanessa L. Bryant

**Affiliations:** Immunology Division, Walter and Eliza Hall Institute of Medical Research, Parkville, Victoria, Australia; Department of Medical Biology, University of Melbourne, Parkville, Victoria, Australia; Snow Centre for Immune Health, Parkville, Victoria, Australia; Department of Clinical Immunology & Allergy, The Royal Melbourne Hospital, Parkville, Victoria, Australia; Department of Gastroenterology, The Royal Melbourne Hospital, Parkville, Victoria, Australia

## Abstract

T cells integrate signals from antigen and co-stimulatory receptors to calibrate the strength and quality of their responses. This signal integration is influenced by genetic background, which can modulate thresholds for immune tolerance and strength of responses to threat. Celiac disease (CeD) is an autoimmune disorder driven by well-defined genetic risk and characterised by immune dysregulation in response to dietary gluten. However, whether sensitivity or response differences in naive T cell programming contributes to disease susceptibility is not known. To investigate such variation, we developed a sensitive quantitative platform, ‘the momentum assay’, which combines standardised, T cell activation with subsequent stimulus withdrawal, enabling measurement of T cell proliferation and survival over time. This assay is integrated with the Cyton2 mathematical model to infer underlying cellular timer programs from population-level dynamics.

We applied this method to assess whether naïve T cells from individuals with celiac disease (CeD) exhibited altered responses compared to healthy donors (HDs). We found that CD4^+^ but not CD8^+^ T cells from CeD patients showed a hypo-proliferative response following stimulation, associated with impaired secretion of the proliferative and pro-survival cytokine IL-2. Moreover, surface expression of early activation marker, CD69 remained elevated for longer on CD4^+^ T cells from CeD donors after stimulus withdrawal, suggesting prolonged activation and subtle alterations in regulatory feedback mechanisms. These findings reveal previously unrecognised quantitative alterations in naïve T cell programming in CeD and underscore the utility of model-based analytical frameworks for detecting subtle functional perturbations in complex immune-mediated diseases.

## Introduction

Naïve T cells are the cornerstone of adaptive immunity, comprising a pool of antigen-inexperienced lymphocytes poised to respond to novel challenges. These cells are guided by germline-encoded genetic programs and environmental signals to make decisions governing rates of activation, proliferation and differentiation upon encountering specific antigen. Because they have not yet undergone antigen-driven clonal expansion or effector commitment, naïve T cells retain molecular signatures that reflect their inherited immune programming. As such they have potential to reveal intrinsic immune dysregulation underlying complex diseases where the causes of inappropriate immune activation are multifactorial, and the molecular determinants of disease susceptibility remain poorly defined.

Celiac disease (CeD) is a chronic immune-mediated enteropathy precipitated by exposure to dietary gluten in genetically susceptible individuals (1). It is characterised by an acquired adaptive immune response to post-translationally modified (deamidated) gluten peptides (2) presented by HLA-DQ2.5, 2.2 and/or HLA-DQ8 molecules to CD4^+^ T cells (3). Once established, this interaction drives rapid release of interleukin-2 (IL-2) and acute gastrointestinal symptoms within two hours of gluten exposure, followed by specific expansion of gluten-reactive CD4^+^ T cells (4) and a broader, non-specific activation of gut-homing CD8^+^ T cells within one week (5). Sustained gluten exposure results in the characteristic histological features of CeD in the small intestine, including diagnostic villous atrophy and crypt hyperplasia (6). The gluten-specific CD4^+^ T cell pool persist as long-lived effector memory clonotypes in both the gut and circulation, reactivating upon re-exposure to gluten (7). CD8^+^ intraepithelial lymphocytes also contribute to tissue pathology, acquiring cytolytic NK-like features in proinflammatory environments and are implicated in mediating the enteropathy of CeD (2).

Genetic studies have identified the strongest CeD risk is conferred by specific HLA alleles encoding disease-permissive MHC Class II molecules HLA-DQ2.5 and HLA-DQ8 (8). Beyond HLA, over 40 non-HLA loci, many involved in T-cell signalling and regulation (e.g. *IL2, PTPN2, CD28, STAT4*) also contribute to CeD susceptibility, pointing to a broader dysfunction in T-cell responses (9–12). Additionally, transcriptomic analyses of gluten-specific CD4^+^ T cells from individuals with CeD further reveal complex activation-associated dysregulation, suggesting that intrinsic abnormalities in T-cell function may contribute to disease risk and progression (13).

To date, mechanistic studies have primarily focused on gluten-reactive memory CD4^+^ T cells either following gluten challenge or stimulated ex vivo. While informative, these approaches do not address whether polyclonal naïve T cells, prior to antigen exposure, are intrinsically dysregulated in CeD. Given the polygenic nature of the disease, we hypothesised that subtle, inherited differences in naïve CD4 and CD8 T cell compartments may shape immune responsiveness and influence disease trajectory. Investigating naïve T-cell responses to controlled polyclonal stimuli provides an opportunity to interrogate fundamental immune programming without the confounding effects of memory formation or environmental exposure.

CeD presents a particularly tractable model to interrogate intrinsic T-cell dysfunction. Unlike many autoimmune disorders managed with immunosuppressive medications, CeD is treated solely through dietary antigen exclusion via strict adherence to a gluten-free (GF) diet. This offers a unique opportunity to dissect intrinsic differences in fundamental T-cell response processes in an autoimmune setting without confounding effects of immunosuppression, which can alter cell survival and perturb signalling and response thresholds (14). Moreover, individuals with treated CeD (on a GF diet) and those with active, newly diagnosed disease can be studied in parallel, enabling direct comparisons across antigen exposure and inflammatory states.

To explore whether naïve T cells in CeD exhibit intrinsic functional abnormalities, we developed a novel *T cell momentum assay* that enables detailed analysis of lymphocyte fate decisions under defined, controlled conditions. In this assay, polyclonal naïve T cells are subjected to strong, non-specific stimulation for 42 hours, after which the stimulus is withdrawn. The subsequent short burst of proliferation, survival and phenotype changes are monitored, as a measure of how effectively T cells integrate and respond to activation signals within a defined temporal window. Importantly, the assay controls for changing levels of IL-2, a major confounding variable in conventional T cell in vitro assays, thereby enabling isolation of intrinsic differences in T cell responsiveness. The response conforms to the Cyton2 model (15) allowing quantification of key parameters such as time to first cell division, division destiny (DD) and time to death, while capturing the stochastic nature of lymphocyte fate decisions through probability distribution functions (16–21).

While previous studies using murine *in vitro* systems and division-tracking dyes such as Cell Trace Violet (CTV) have provided key insights into the molecular regulation of T-cell proliferation and survival (22–26), the T cell momentum assay offers a novel application of these principles to human naïve T cells. Thus, we measure how well T cells accumulate signals in a short period by measuring their proliferation and survival ‘momentum’ after stimulus removal. Using this approach, we compared proliferative and survival responses of polyclonal naive CD4^+^ and CD8^+^ T cells from individuals with CeD and healthy donors (HDs). We identified significant differences in the naïve CD4^+^ T cell compartment of individuals with CeD, including altered IL-2 secretion and CD69 expression. These findings suggest that naive CD4^+^ T cells in individuals with CeD are functionally distinct at baseline, implicating inherited molecular programming in predisposing the immune system towards dysregulated activation prior to antigen encounter.

## Results

### Development of the human naïve T cell momentum assay

We first aimed to establish a human naïve T cell assay to quantify proliferation, division and survival of activated naïve CD4^+^ and CD8^+^ T cells from HD and CeD individuals. We recognised that conventional T cell stimulation assays that allow endogenous IL-2 secretion are highly sensitive to starting cell density making quantitative comparisons difficult. To this end, we developed a novel momentum assay to probe the programmed proliferative potential accumulated by human naïve T cells during a 42-hour activation period that included anti-CD3, anti-CD28 and IL-2, followed by assessment of their proliferation, division and survival after stimulus withdrawal (Figure 1A). Naïve CD4^+^ and CD8^+^ T cells were isolated and labelled with CTV. Post-purity assessment was ∼95-96% (Supplemental Figure 1A). CTV-labelled cells were cultured in the presence of activating anti-CD3/anti-CD28 coated Dynabeads and of recombinant human IL-2 (rhuIL-2) for 42 hours to ensure maximal activation. T-cell responses, including survival and division during the stimulation period (0-42 hours) were measured prior to stimulus removal.

**Figure 1.**
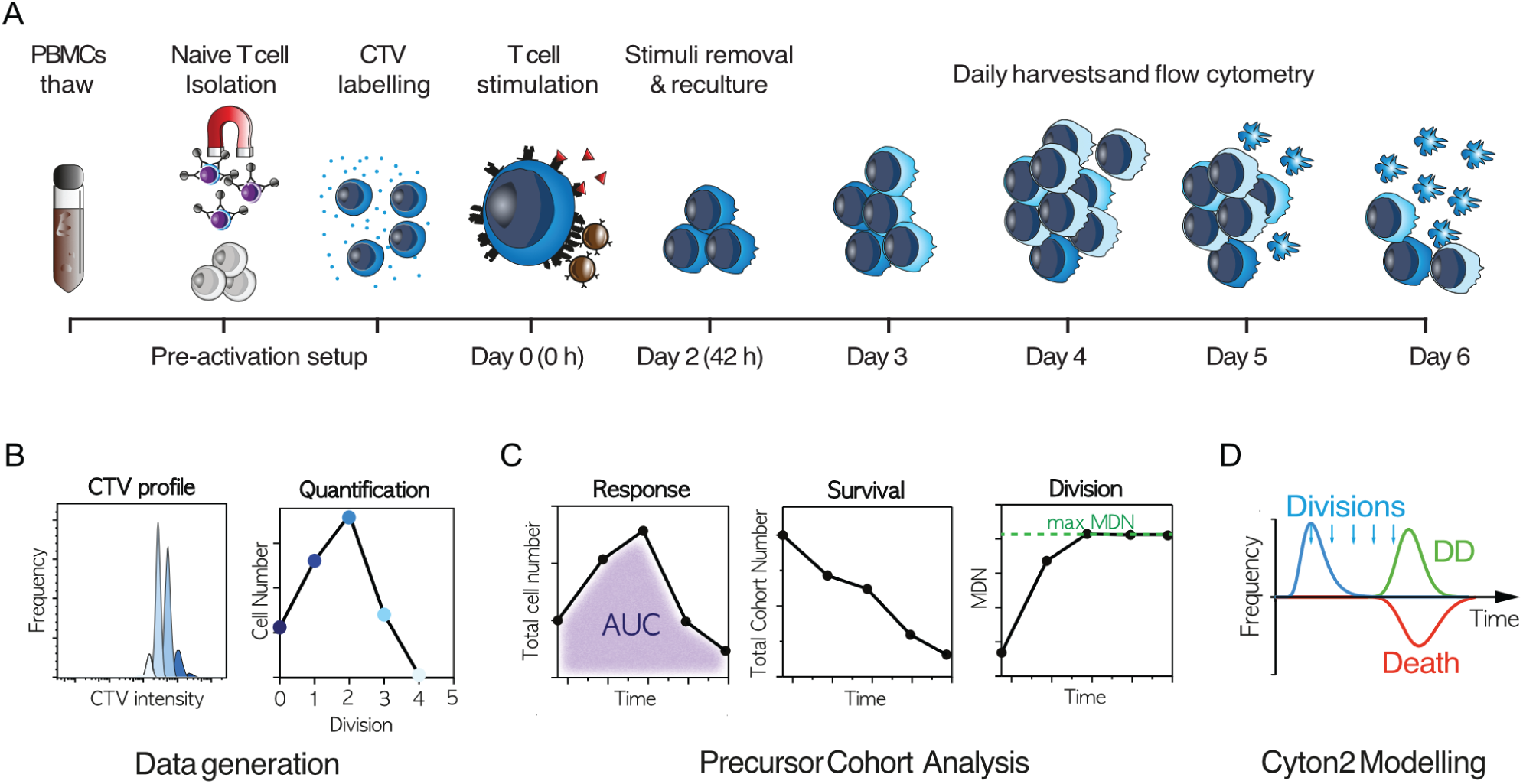
A functional assay to assess division momentum and fate programming in human naïve T cells. **(A)** Overview of the human naïve T cell momentum assay. Cryopreserved PBMCs from HD and CeD individuals were thawed and naïve CD4^+^ or CD8^+^ T cells isolated via negative isolation. Cells were labelled with division-tracking dye, Cell trace violet (CTV), and stimulated with anti−CD3/CD28 Dynabeads and 100 U/mL rhuIL-2. After 42 hours, activated T cells were washed (and beads magnetically detached) to remove activation signals and recultured for an additional 4 days in media alone or in the presence of blocking IL-2 conditions (anti-IL-2/IL-2Rα) with or without rhuIL-2 (31.6 U/ml). Cells were harvested daily and analysed using flow cytometry. **(B)** Quantitative metrics extracted from the assay. Proliferation was measured by analysing CTV dilution profiles and calculating precursor cohort numbers within each division generation. **(C)** Precursor cohort analysis enabled quantification: Total cell number to determine overall response magnitude; total cohort number, reflecting survival of the initial input population over time; and mean division number (MDN), for quantification of average division of cell population. Area under the curve (AUC) of total cell numbers over time was used to derive a single proliferative value. Maximum MDN reflective was used as a summary of overall division burst. **(D)** Cell fate timer distributions (time to first division, time to death and time to division destiny, DD) were estimated using the Cyton2 mathematical model, with each modelled as a lognormally distributed random variable.

After 42 hours, activation signals were withdrawn, and cells were re-cultured under three conditions: fresh media alone; media supplemented with anti-IL-2/anti-IL-2Rα antibodies to neutralise endogenous IL-2; or media containing anti-IL-2/IL-2Rα plus exogenous rhuIL-2. The latter condition provided a survival advantage compared to IL-2 blocking conditions alone without promoting further division (Supplementary Figure 2), allowing for accurate quantification of mean division number (MDN), which would otherwise decline due to preferential death of cells in later division generations. Proliferation kinetics were tracked for an additional four days, for a total of 6 days from initial stimulation.

CTV profiles were acquired daily using flow cytometry (Supplemental Figure 1B), and cell counts per division were quantified (Figure 1B). Summary statistics such as total cell numbers, total cohort numbers, and MDN over time were derived using the precursor cohort method (16, 25, 27), which provides interpretable metrics reflecting response magnitude, population survival and division progression, respectively (Figure 1C). The area under the curve (AUC) of normalised total cell counts was also computed as an aggregate measure of proliferative output. To gain mechanistic insight into the cell population dynamics underpinning these responses, we employed the Cyton2 model, which estimates parameters including three stochastic variables: time to first division, time to division destiny (DD), time to cell death, alongside a constant for subsequent division intervals (Figure 1D). The model assumes lognormal distributions for these variables and provided an excellent fit to the experimental data.

To ensure robustness of the momentum assay, we titrated cell density, stimuli and reagents to confirm consistency across experimental setups. Comparable proliferative responses were observed across varying cell densities of cells (both at the initial culture and after re-culture) and with different Dynabead ratios, indicating minimal impact from small deviations from our initial starting numbers would have little effect on results (Supplemental Figure 2A-C). Addition of anti-IL-2 and anti-IL2Rα with rhuIL-2 (31.6 U/ml) enhanced cell survival without affecting division (Supplemental Figure 2D-E), as cells in this condition did not show the selective death of later-generation cells observed in anti-IL-2/IL-2Rα-only cultures (Supplemental Figure 2F-H). Furthermore, expression levels of the division regulator, Myc (22) declined to baseline at the same rate in both cultures (Supplemental Figure 2I). These findings provided confidence in our setup to assess T-cell responses from CeD donors.

We next determined whether sex or age influenced naïve CD4^+^ and CD8^+^ T-cell responses in HD samples. No significant sex-specific differences (Supplemental Figure 3) or age-related correlations (Supplemental Figure 4) were observed across any of the proliferative parameters assessed for both CD4^+^ and CD8^+^ T cells. Having validated the momentum assay in healthy donors, we next applied it to our CeD cohort in parallel with HD samples to evaluate the health of naïve T-cell responses and to explore biologically meaningful differences.

### CD8^+^ T cells from CeD individuals exhibit enhanced early survival during activation

Upon stimulation, cell signalling and genetic reprogramming events transform quiescent, naïve T cells into activated and proliferating T cells. To assess early activation responses, purified CTV-labelled naive CD4^+^ or CD8^+^ T cells from HD and CeD individuals were stimulated for 42 hours with anti-CD3/anti-CD28 Dynabeads in the presence of 100 U/mL rhuIL-2. Both CD4^+^ and CD8^+^ T cells had begun to divide at this timepoint (Figure 2A).

**Figure 2.**
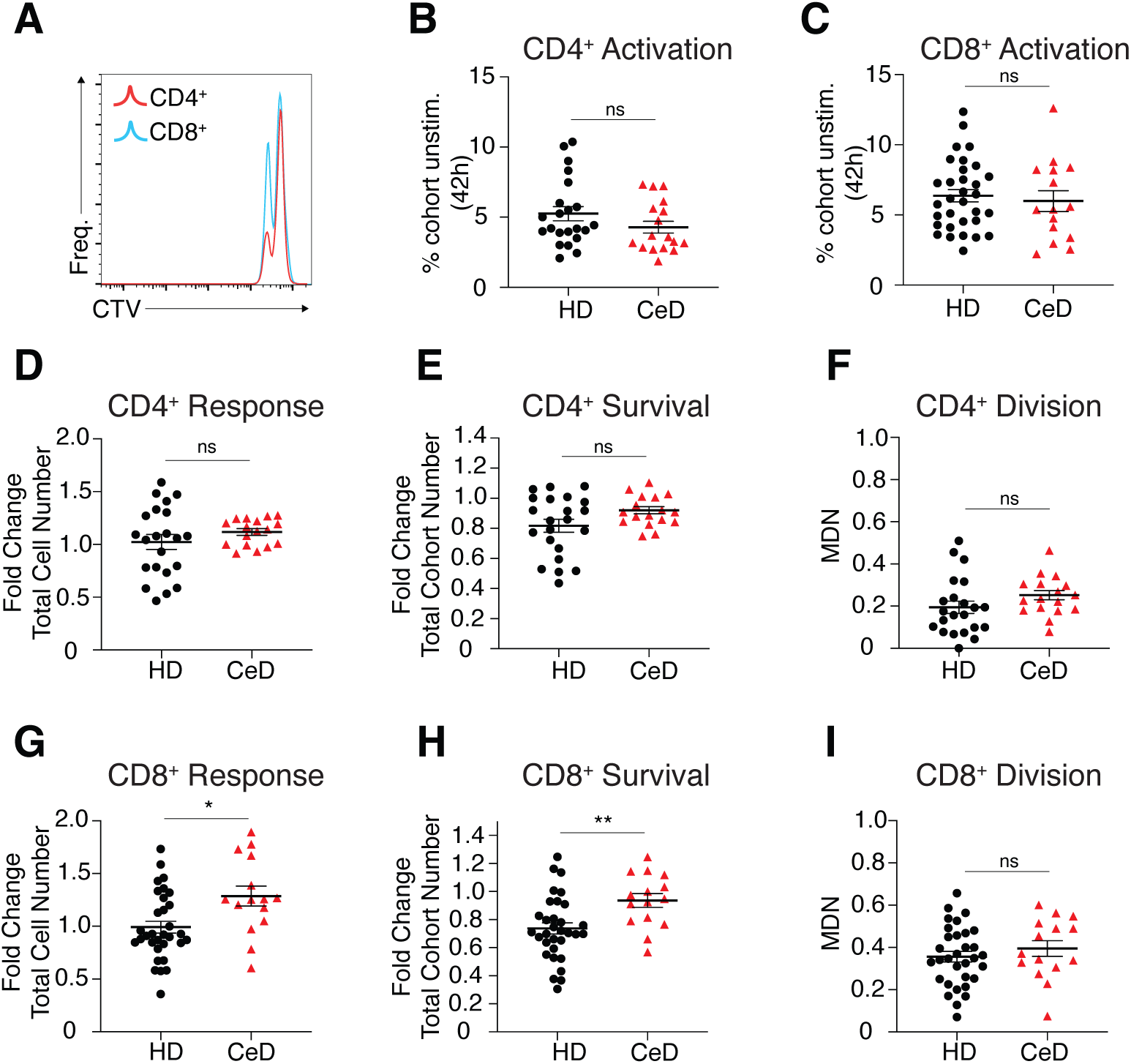
Enhanced early survival of naive CD8^+^ T cells from CeD donors prior to stimulation removal. Naïve CD4^+^ and CD8^+^ T cells from healthy donors (HD) and individuals with Celiac Disease (CeD) were stimulated using the momentum assay: stimulated for 42 hours with anti−CD3/CD28 Dynabeads and 100 U/mL rhuIL-2. **(A)** Representative CTV proliferation profiles from activated naïve CD4^+^ and CD8^+^ T cells after 42 hours. **(B, C)** Frequencies of CD4^+^ (B) and CD8^+^ (C) T cells were measured based on CTV dilution and size as an estimate of T-cell activation. **(D-F)** Quantification of 42-hour CD4^+^ T-cell response from HD and CeD donors: (D) total cell number (overall response), (E) total cohort number (survival), and (F) mean division number (MDN; proliferation). **(G-I)** Quantification of 42-hour CD8^+^ T-cell response: (G) total cell number, (H) total cohort number and (I) MDN. Data are presented as fold-change relative to matched unstimulated controls. Mean ± SEM for HD and CeD groups. Sample sizes: CD4^+^ T cells (HD = 22, CeD = 17); CD8^+^ T cells (HD = 32, CeD = 15). Comparisons performed using unpaired t-test with Welch’s correction. *indicates p<0.05; ** indicates *p*<0.01.

To compare levels of activation, we quantified the proportion of unstimulated naïve T cells remaining in culture. Small, undivided cells, morphologically resembling naive unstimulated cells, were gated using FSC-A vs CTV plots (Supplemental Figure 1B) and expressed as a proportion of the total cohort. Both HD and CeD samples showed robust activation, with only ∼5% CD4^+^ (mean ±SEM) and ∼6% CD8^+^ (mean ±SEM) T cells remaining unstimulated at 42 hours (Figure 2B, C). No significant differences were observed in total cell numbers, total cohort numbers (indicative of cell survival) or MDN between HD and CeD samples for CD4^+^ T cells (Figure 2 D-F). In contrast, CD8^+^ T cells from CeD donors showed increased total cell and cohort numbers compared to HD, while MDN remained comparable (Figure 2G-I) indicting enhanced survival rather than accelerated division.

To explore whether active inflammatory status in CeD influenced early CD4^+^ responses, we stratified CeD donors based on disease status: those following a gluten-free (GF) diet and those with newly diagnosed active CeD still consuming gluten (Supplemental Figure 5A). CD4^+^ T cells from donors with active CeD exhibited a small, but significant increase in total cell numbers and survival compared to treated CeD donors and healthy controls, however; no difference was observed in activation or division. These findings suggest that disease state had a small underlying effect on early cell survival within CD4^+^ T cells from active but not treated CeD individuals. Together, early kinetic analysis confirms that naive CD4^+^ and CD8^+^ T cells from both HD and CeD donors undergo robust activation within 42 hours. However, CD8^+^ T cells from CeD donors, and to a lesser extent CD4^+^ T cells from active CeD cases, display slightly enhanced survival during this early phase.

### Hypo-proliferative naïve CD4^+^ T-cell responses in CeD donors

We next assessed the proliferative momentum of naive T cells from HD and CeD donors following stimuli withdrawal. After 42 hours of activation, anti-CD3/CD28 activator beads were removed, and activated T cells were washed and recultured into one of three conditions: media alone, IL-2 blocking condition (media with anti-IL-2/IL-2Rα blocking antibodies to block endogenous IL-2), or IL-2 blocking plus 31.6 U/mL exogenous rhuIL-2. The increased survival without affecting cell division observed by the addition of rhuIL-2 in the presence of IL-2 blockade (Supplemental Figure 2D-I), enabled analysis of division cessation independent of survival bias in later generations.

Proliferation, survival and division kinetics were measured for six days post-initial stimulation. Notably, in the media alone condition, naïve CD4^+^ T cells from CeD showed significantly lower total cell numbers compared to HD from day 4 onwards (Figure 3A). This hypo-proliferative phenotype was linked to reduced total cohort numbers over time, indicating impaired cell survival. Division kinetics, including division rate and maximum division generation reached, were comparable, although a small reduction in MDN was observed at days 5 and 6 in CeD samples, consistent with preferential cell death in later-generation cells. No differences were observed between active and treated CeD groups (Supplemental Figure 5B). Furthermore, unstimulated naïve CD4^+^ and CD8^+^ T cells from both HD and CeD donors displayed similar rates of cell death (Supplemental Figure 5C, D), indicating that survival differences were specifically in response to stimulation.

**Figure 3.**
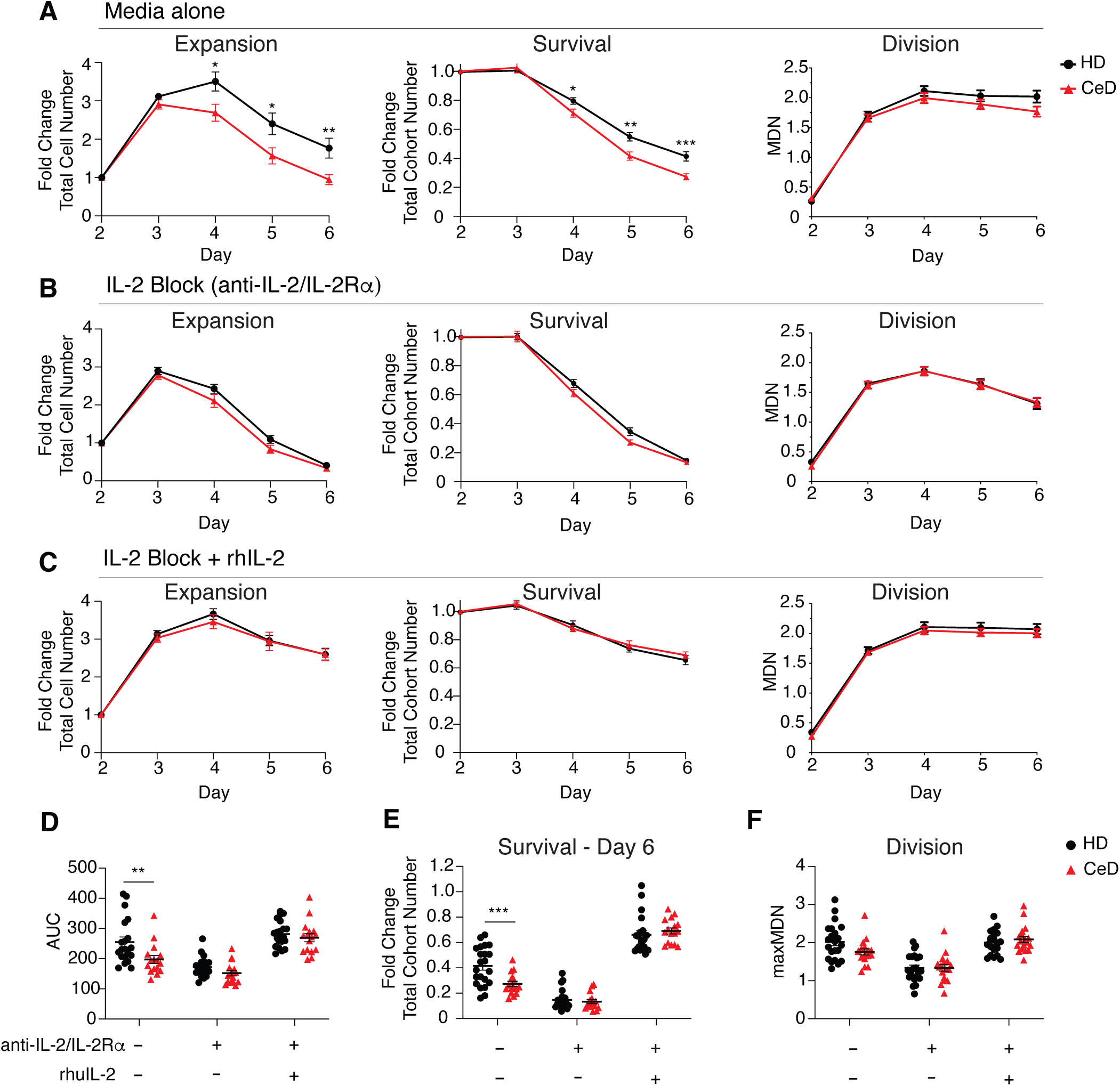
Hypo-proliferation of naïve CD4^+^ T cells from CeD donors is IL-2 dependent. Naïve CD4^+^ T cells from HD and CeD donors were stimulated for 42 hours with anti−CD3/CD28 Dynabeads and 100 U/mL rhuIL-2, followed by stimulus removal. Cells were cultured for an additional 4 days in media alone or with IL-2 blocking conditions and harvested daily for flow cytometric analysis. **(A)** CD4^+^ T cells recultured in media alone: fold change total in (left) total cell number, (middle) total cohort number (survival) and (right) MDN (proliferation). **(B)** CD4+ T cells cultured with anti-IL-2/IL-2Rα antibodies: (left) total cell number, (middle) total cohort number and (right) MDN. **(C)** CD4+ T cells cultured with anti-IL-2/IL-2Rα antibodies plus exogenous rhuIL-2: (left) total cell number, (middle) cohort number and (right) MDN. **(D-F)** Summary measures across conditions (D): area under the curve (AUC) for total cell number, (E) fold-change in total cohort number at day 6, and (F) maximum MDN. Data shown as mean ± SEM; sample sizes: HD = 22, CeD = 16. Comparisons were made using unpaired t-test with Welch’s correction. ** indicates p<0.01; *** indicates p<0.001.

To investigate whether differences in IL-2 produced by activated T cells contributed to impaired CD4^+^ T cell survival in CeD, we blocked endogenous IL-2 using ant anti-IL-2/IL-2Rα antibodies. This eliminated survival differences and normalised expansion kinetics between HD and CeD groups (Figure 3B). Moreover, addition of exogenous rhuIL-2 to these cultures enhanced survival equally in both groups (Figure 3C), further supporting a defect in autocrine IL-2 mediated survival signalling in CeD-derived CD4^+^ T cells.

AUC analysis of total cell numbers revealed significantly lower responses in CeD samples under media-alone conditions, but not when IL-2 signalling was neutralised or rescued (Figure 3D). Similarly, total cohort number at day 6 were significantly reduced under media-alone conditions (Figure 3E). MDN remained comparable between HD and CeD samples across all conditions (Figure 3F), providing further evidence that the proliferative defect was due to impaired survival rather than defective division. In contrast, CD8^+^ T cells from CeD donors did not exhibit the same IL-2-dependent differences in expansion kinetics (Supplemental Figure 6). These findings indicate that naïve CD4^+^ T cells from CeD donors, exhibit dysregulated response upon activation, characterised by hypo-proliferation driven by impaired IL-2 dependent survival.

### Cyton2 modelling reveals early death timers in CD4^+^ T-cell responses from CeD donors

To gain mechanistic insight into the hypo-proliferative phenotype observed in CD4^+^ T cells from CeD donors, we applied the Cyton2 model to division data from cells cultured in media alone (from Figure 3A). This model allows estimation of stochastic cellular fate parameters-specifically, the medians and log-variances of the distributions for timers of first division (T^0^_div_), death (T_die_) and division destiny (T_dd_) – as well as the average time between subsequent divisions (parameter *b*). Best-fit model and parameter values and 95% confidence intervals were obtained using least-squares fitting and bootstrap methods, respectively (Figure 4A). The model was fitted to observed total cell numbers per generation over time across both HD and CeD cohorts (Figure 4B, see https://github.com/hodgkinlab/CeDmomentum2025). Empirical cumulative distribution functions (eCDFs) were generated for each fate timer using cohort-specific median (*m*) values. This analysis revealed that CD4^+^ T cells from CeD donors exhibited significantly earlier death times (m_die_) compared to HD, while time to first division (m^0^_div_) and division destiny times (m_dd_) were comparable between groups (Figure 4C). To test the significance of these differences, we performed hypothesis testing using a non-parametric permutation test. Here, the observed difference in medians between HD and CeD cohorts was compared to a null distribution generated from 10^7^ random permutations. This confirmed that the earlier death time in CeD (time ± CI) was statistically significant, while differences in m^0^_div_ and m_dd_, were not (Figure 4D).

**Figure 4.**
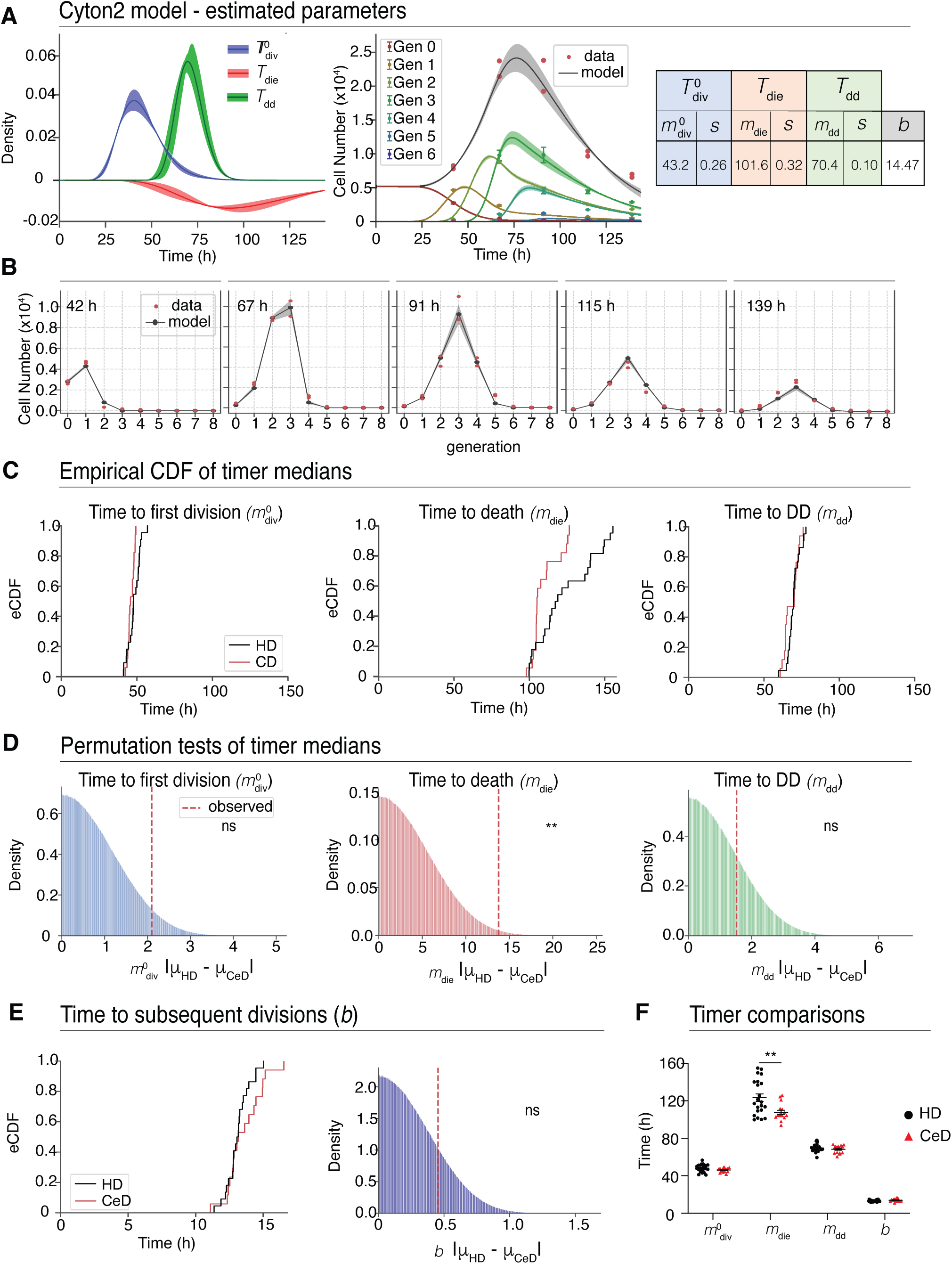
Cyton2 modelling reveals earlier cell death in CD4^+^ responses from CeD donors. Cyton modelling was applied to CD4^+^ T cell momentum assay data to estimate underlying cell fate timers. **(A)** Example best-fitted Cyton2 parameters showing lognormal distributions (±95% CI) for time to first division (T^0^_div_), time to die (T_die_) and time to DD (T_dd_; left panel). Model extrapolation of total cell numbers in generations 0-6 over time (middle). Estimated median (*m*) and log-variance (*s*) for T^0^_div_, T_die_, T_dd_ and a constant for subsequent division time (*b*) for represented donor sample (right). **(B)** Model fit to observed cell number per generation for each time-point. **(C)** Empirical cumulative distribution functions (eCDFs) comparing HD and CeD: median time to first division (*m*^0^_div_), median time to die (*m*_die_) and median time to DD (*m*_dd_), as indicated. **(D)** Permutations testing comparing observed differences between average HD and CeD medians (red dashed line) for: *m*^0^_div_, *m*_die_, and *m*_dd_. **(E)** eCDF comparing subsequent division constant (*b*) between HD and CeD cohorts. Permutation test comparing average subsequent division (*b*) between HD (black) and CeD (red) cohorts. **(F)** Summary comparison of medians model parameters (*m*^0^_div_, *m*_die_, *m*_dd_) and subsequent division constant (*b*) between groups. Data are presented as means of timer medians and *b* ± SEM; (HD = 22 CeD = 16). Comparisons used unpaired t-tests with Welch’s correction.

We further assessed the average time for subsequent divisions (*b*) and found no significant differences between cohorts (Figure 4E). This was confirmed using t-tests on cohort level medians for each timer and subsequent division, which again demonstrated a significant reduction in m_die_ for CeD donor samples, but no differences in m^0^_div,_ m_dd_ or *b* compared to HD (Figure 4F). Finally, we examined whether IL-2 signalling accounted for any observed differences in cellular timer distributions. When Cyton2 modelling was applied to cultures in which IL-2 was neutralised or supplemented (as in Figure 3), no differences in timer medians were observed between HD and CeD cohorts for either CD4^+^ or CD8^+^ T-cell responses (Supplemental Figure 7). Together, Cyton2 modelling provides quantitative evidence that the impaired proliferative capacity of CD4^+^ T cells from CeD donors is driven by altered cell fate dynamics in cultures lacking control of endogenously produced IL-2.

### Activated naïve CD4^+^ T cells from CeD donors secrete reduced levels of IL-2, contributing to hypoactive responses

Given that impaired CD4^+^ T cell survival was only observed in cultures where the effects of endogenously produced IL-2 were not controlled, we next investigated whether changes in IL-2 secretion or expression of the high-affinity IL-2Rα (CD25) could explain the hypo-proliferative responses seen in CeD donors. To assess this, we first measured IL-2 concentrations and IL-2Rα (CD25) expression on day 3 of the momentum assay, following removal of exogenous rhuIL-2. IL-2 levels were significantly lower in cultures from CeD donors compared to HD, while CD25 expression was comparable between groups (Figure 5A). IL-2 secretion levels in culture supernatants significantly correlated (HD p<0.01, CeD p<0.05) with AUC for total cell numbers (Figure 5B). Regression analysis comparing estimated slope and elevation indicated no significant difference between CeD and HD cohorts (Figure 5B), suggesting that IL-2 signalling was intact and that both groups responded similarly to equivalent concentrations.

**Figure 5.**
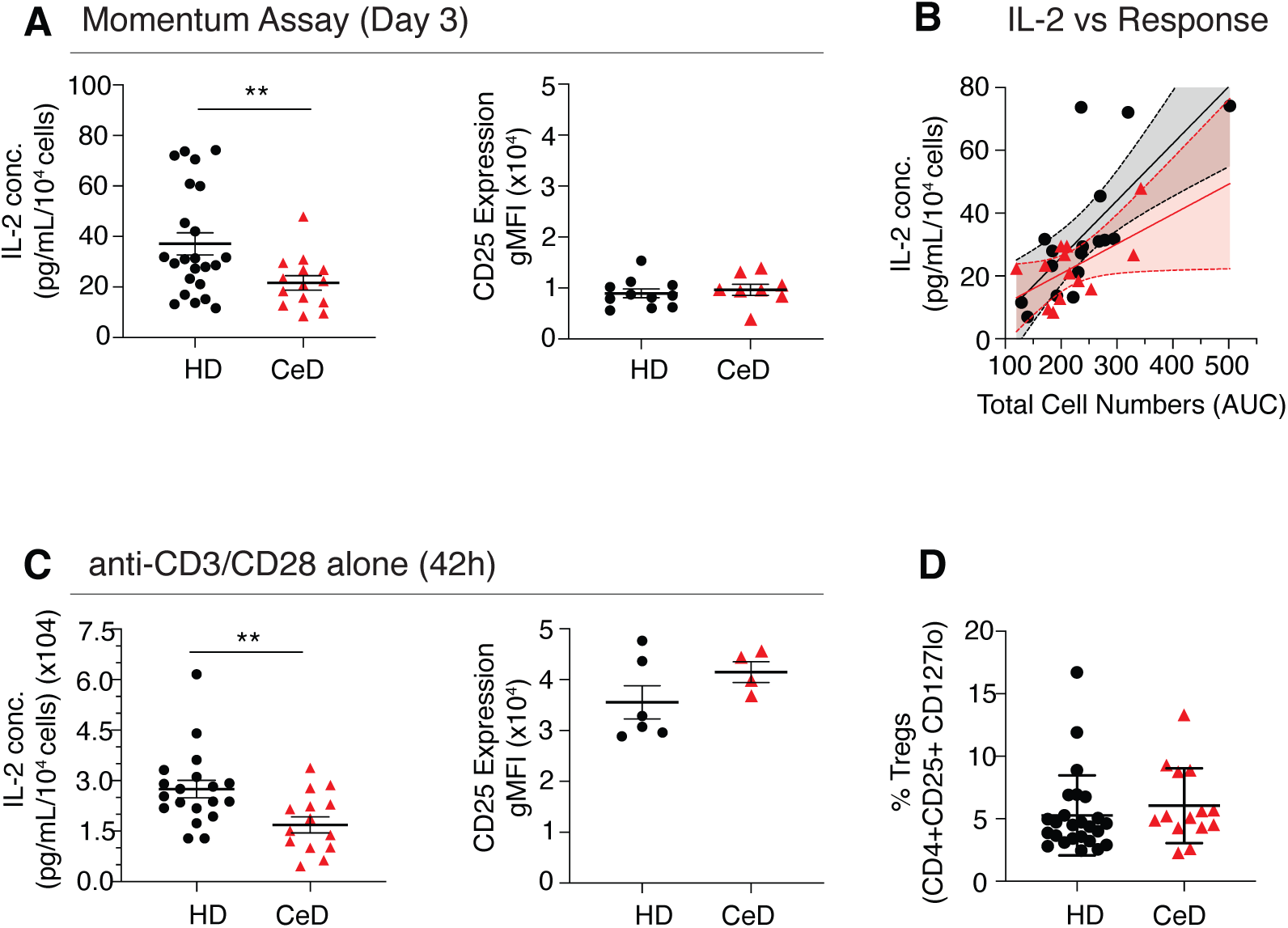
Reduced IL-2 secretion contributes to hypo-proliferative responses of activated naïve CD4^+^ T cells from CeD donors. Naïve CD4+ T cells from HD and individuals with CeD were stimulated using the momentum assay and IL-2 production and IL-2Rα expression were assessed following activation. **(A)** IL-2 concentration was measured in supernatants collected at day 3 of the momentum assay from CD4+ T cell cultures originally stimulated with αCD3/CD28 Dynabeads without addition of IL-2 for 42 h (HD = 23, CeD = 14). Surface expression of CD25, the high affinity IL-2 receptor alpha chain, was quantified by flow cytometry (HD = 6, CeD = 4). **(B)** Correlation between IL-2 secretion (pg/ml/10^4^ cells) and cumulative proliferation (total cell number AUC) in momentum assay across HD and CeD donors, with linear regression equations: HD, Y = 0.18X –10.38; CeD, Y = 0.11X – 2.82. **(C)** To assess early IL-2 production, supernatants and cells were collected at 42 h from cultures containing anti-CD3/CD28 Dynabeads alone. IL-2 concentration (left; HD = 19, CeD = 14) and CD25 expression (right; HD = 11, CeD = 8) on CD4+ T cells were quantified. **(D)** The frequency of circulating Tregs (CD4^+^ CD25^+^ CD127^-^) was measured ex vivo in PBMCs (HD = 25; CeD = 14) as an overall percentage of T cells. Data are presented as mean values ± SEM. Statistical comparisons were performed using unpaired t-test with Welch’s correction. Linear regression was used to compare slopes and intercepts in (C). ** indicates *p*<0.01.

To further validate these findings, naïve CD4^+^ T cells were activated with anti-CD3/CD28 without addition of rhuIL-2 for 42 hours. Again, endogenous IL-2 secretion was significantly reduced in CeD donors compared to HD, while CD25 expression remained unchanged (Figure 5C). In contrast, CD8^+^ T cells showed no differences in IL-2 secretion or CD25 expression between HD and CeD cohorts (Supplemental Figure 8A-D), consistent with similar proliferation profiles. Stratification by disease state (treated vs active CeD) revealed no significant differences (Supplemental Figure 8E, F). Together, these data demonstrate that reduced IL-2 secretion by activated naïve CD4^+^ T cells in CeD underpins their impaired proliferative capacity.

We next addressed whether this reduction in IL-2 might also impact T regulatory (Treg) cell frequencies, as these cells require IL-2 for survival and homeostasis, as a potential driver for disease in CD. The proportion of circulating Tregs (CD3^+^CD4^+^CD25^+^CD127^-^ cells) was determined in PBMCs from CeD (n=14) and HD (n=25) (Supplemental Figure 9A). Similar proportions of Tregs in the CD4^+^ T cell pool were observed between HD and CeD cohorts (Figure 5D), indicating no overt reduction in circulating Tregs.

### Persistent CD69 expression following stimulus withdrawal in activated naïve CD4^+^ T cells from CeD donors

The observed defect in IL-2 secretion suggests dysregulated integration of TCR and/or CD28 signalling in naïve CD4^+^ T cells from CeD donors. To explore this further, we examined expression of CD69, an early activation marker and downstream target of TCR signalling, as a potential readout of signal processing fidelity. CTV-labelled CD4^+^ T cells were stimulated with anti-CD3/CD28 beads and rhuIL-2 for 42 hours as previously, after which the stimuli was washed off and CD69 expression was measured over time and by division (Figure 6A; B, Supplemental Figure 9B). Interestingly, while no differences were observed immediately after stimuli removal, CD69^+^ cell frequencies were significantly higher in CeD donors 24 hours later (day 3), but this difference resolved at subsequent timepoints. Notably, this transient elevated expression was independent of IL-2 availability in these cultures (Figure 6C).

**Figure 6.**
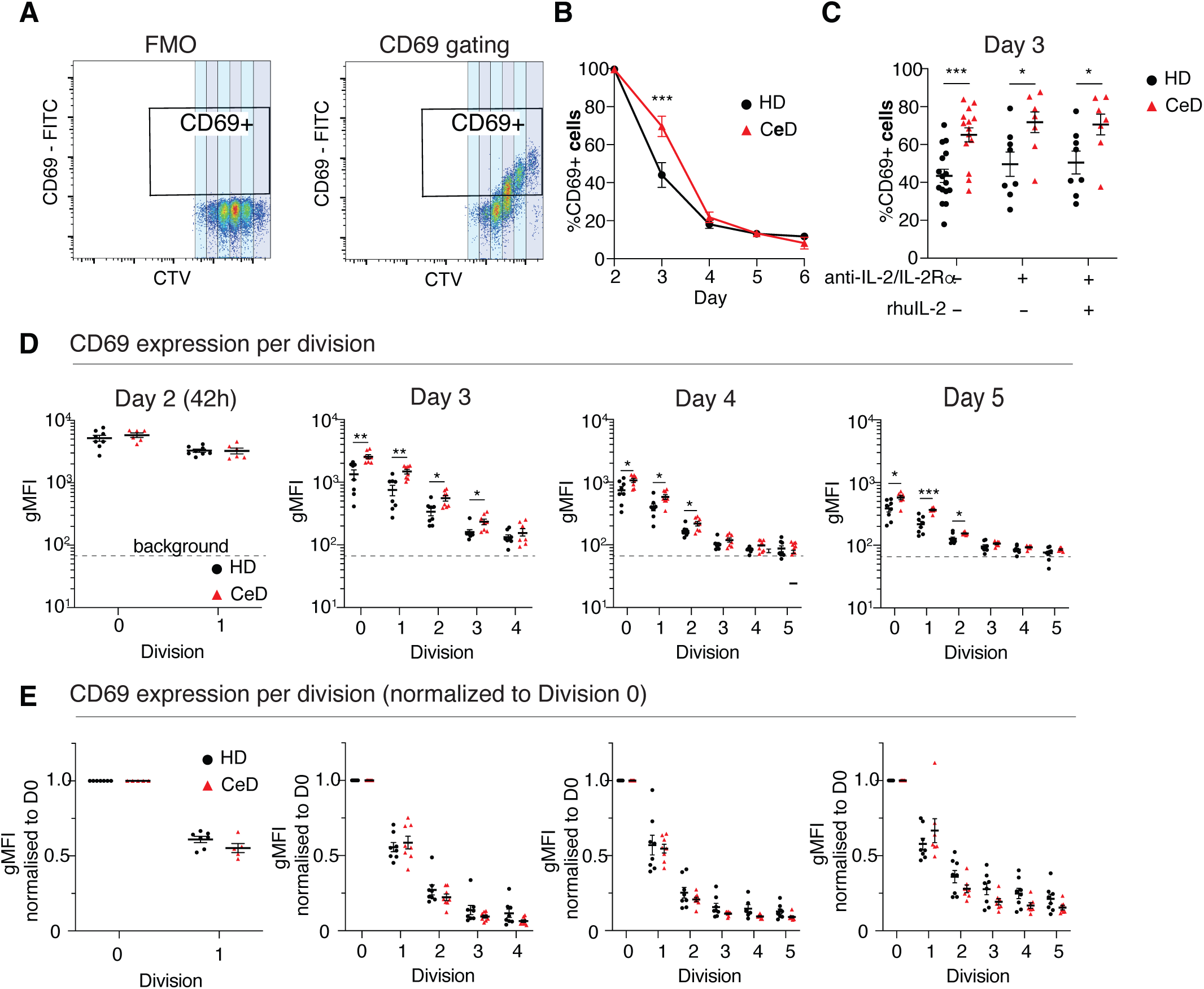
CD69 remains elevated after stimuli removal then declines at a normal rate in naïve CD4^+^ T cell responses from CeD donors. Surface expression of early T-cell activation marker CD69 was measured on CD4^+^ T cells from HD and CeD individuals during the momentum assay. CD69+ cells and division-based kinetics were determined as shown using **(A)** FMO control and gating on CD69 positive cells. **(B)** Percentage of CD69^+^ (% total CD4^+^ T cells over time following activation and stimulus withdrawal (HD, N=8; CeD, N=8). **(C)** CD69 expression was quantified at day 3 post activation in cells cultures under three conditions: media alone (HD, N=16; CeD, N=14), anti-IL-2/IL-2Rα blockade (HD, N=8; CeD, N=7), and anti-IL-2/IL-2Rα blockade with exogenous rhuIL-2 supplementation (HD, N=8; CeD, N=7). **(D)** CD69 gMFI was plotted against division number to assess activation kinetics in dividing cells. **(E)** Fold change CD69 gMFI relative to division 0 across divisions is shown for day 2, 3 and 4 post activation, as indicated (HD and CeD, N=8 each). Data are presented as mean ± SEM. Statistical comparisons were performed using unpaired t-tests with Welch’s correction. * indicates *p*<0.05; ** indicates p<0.01; *** indicates p<0.001.

When stratified by division, elevated CD69 expression was observed within early divisions (generations 0 – 3 on day 3, 0 - 2 on days 4 and 5) in CeD donors (Figure 6D). However, when CD69 gMFI was normalised to undivided (generation 0) levels within each timepoint, no differences were observed between groups (Figure 6E). This suggests that the elevated expression was not driven by division-linked mechanisms but rather reflects delayed CD69 downregulation. Other activation markers, including CD62L and CD127, were similar between HD and CeD donors (Supplemental Figure 9C). Within CD8^+^ T cells, CD69 levels were similar between HD and CeD cohorts (Supplemental Figure 9D, left panel). Interestingly, CD69 on CD4^+^ T cells was significantly higher in individuals with active CeD compared to those on a GF diet. However, both groups showed significantly elevated expression relative to HD controls (Supplemental Figure 9D right panel), suggesting that active disease may exacerbate, but is not solely responsible for, this phenomenon. Collectively, these findings reveal that activated naïve CD4^+^ T cells from CeD donors exhibit delayed downregulation of CD69 following stimulus withdrawal, further supporting a model of dysregulated early signal integration and activation in CeD.

## Discussion

Using our novel T cell momentum assay, we have uncovered evidence of functional dysregulation in polyclonal naïve T cells from individuals with celiac disease (CeD), with the most pronounced effects observed in CD4^+^ T cells. While naïve CD8^+^ T cells from CeD donors exhibited a transient early survival advantage at 42 hours post-stimulation, CD4^+^ T cells displayed a marked hypo-proliferative response following withdrawal of exogenous stimuli. Cyton2 modelling revealed that this impaired proliferation momentum was primarily driven by accelerated cell death, with significantly shortened death timers in CD4^+^ T cells from CeD donors. Importantly, other key parameters such as time to first division and DD timers as well as survival of unstimulated cells remained unaffected, indicating that dysregulated cell death is a specific and defining feature of CD4^+^ T cells in CeD. Mechanistically, we identified reduced IL-2 secretion as a key driver of impaired survival in CeD donor-derived CD4^+^ T cells. This observation is notable, as defects in the naïve T cell compartment have not previously been reported in CeD. These findings raise important questions regarding the origin of naïve T-cell dysfunction in CeD and its potential contribution to disease development, progression, and prognosis.

IL-2 is critical for T cell proliferation and survival (17), yet our data demonstrate that reduced IL-2 secretion by CD4^+^ T cells in CeD selectively compromises cell survival with minimal impact on division timing. Interestingly, similar survival phenotypes were observed in cultures treated with anti-IL-2/IL-2Rα blocking antibodies alongside rhuIL-2, suggesting that blockade of high-affinity IL-2Rα (via CD25) may redirect signalling through low affinity IL-2R (comprising IL-2Rγ and β chains). This alternative signalling may preferentially promote pro-survival over pro-proliferative responses. A similar mechanism may occur in CeD-derived CD4^+^ T cells following stimulus withdrawal, where low endogenous IL-2 may favour such non-canonical IL-2 signalling. The molecular basis by which IL-2 signalling is differentially integrated to promote survival over division in this setting remain unclear and warrants further mechanistic investigation. Nonetheless, our data establish that reduced IL-2 secretion by CD4^+^ T cells is a defining and functionally significant feature of naïve CD4^+^ T cells from CeD donors, suggesting dysregulated integration of TCR and costimulatory signals at the molecular level.

We also observed aberrant kinetics of activation marker CD69 expression in CD4^+^ T cells from CeD donors. Although peak CD69 levels were comparable to those in healthy donors, expression remained elevated for a short period following removal of stimulatory signals. As CD69 is a rapidly induced early activation marker typically downregulated upon termination of TCR signalling (28), its sustained elevated expression suggests a delay in the downregulation of proximal TCR signalling pathways in CeD T cells. This may reflect impaired negative regulatory mechanisms that normally constrain T-cell activation dynamics. Although heightened sensitivity to TCR stimulation and increased TCR affinity for self-antigens have been implicated in the pathogenesis of autoimmune disorders (29), our quantitative momentum assay, specifically designed to assess intrinsic TCR responsiveness, did not reveal evidence of TCR hypersensitivity in naïve CD4^+^ T cells from CeD donors.

Together, these findings reveal a previously unrecognised functional defect in the naïve CD4^+^ T cell compartment in CeD, characterised by impaired survival due to reduced IL-2 secretion and delayed deactivation of TCR signalling. This immune phenotype may represent an early functional risk state that contributes to disease susceptibility or progression and has important implications for understanding immune dysregulation underlying CeD, and autoimmune conditions, more broadly.

The proliferative differences observed in CD4^+^ T-cell responses from CeD donors reflect an impaired ability of these cells to integrate stimuli. This dysfunction likely stems from intrinsic defects in the sensing and interpretation of stimulatory inputs, potentially involving abnormal receptor and/or downstream signalling processes. Indeed, TCR signalling via CD3 and costimulatory receptor CD28 converges on pathways that activate key transcription factors driving IL-2 production (30, 31). Aberrant signalling may therefore arise from dysregulated receptor function, defects in downstream signalling proteins, or impaired transcriptional responses - alone or in combination. Supporting this hypothesis, genome-wide association studies (GWAS) have linked CeD risk to variants in genes involved in T-cell activation, including loci within the IL-2 and associated signalling pathways (9, 10, 12). Further, transcriptomic analyses of bulk CD4^+^ T cell populations from CeD patients have similarly revealed differential expression of genes involved in T-cell activation, cytokine signalling, and other critical immuno-molecular pathways (30, 44). It is plausible that this naïve T cell dysfunction in CeD arises early, driven by germline polymorphisms affecting T-cell activation or cytokine production, and this is further shaped by epigenetic modifications influenced by environmental exposures, such as infections or diet. Longitudinal birth cohort studies of genetically at-risk infants will be critical to determine whether these naïve T cell abnormalities are present at birth or emerge later prior to clinical onset of CeD. Predictive tools such as genomic risk scores (32, 33) and disease-specific proteomic signatures in blood (34) both demonstrate a potential for identifying individuals likely to develop CeD, further supporting the hypothesis that genetically-determined immune dysfunction is a likely precursor to overt CeD development. Importantly, a lowered capacity for IL-2 production by naïve CD4^+^ T cells may represent a key mechanistic link in the breakdown of tolerance and disease development. Previous studies have indicated that insufficient IL-2 signalling, whether due to limited cytokine availability or defects in IL-2 receptor components, can drive autoimmune and autoinflammatory outcomes (35–37). The role of Tregs becomes particularly relevant in this context, as they rely on adequate IL-2 for their development, expansion and suppressive function (38). If naïve CD4^+^ T cells in CeD are intrinsically impaired in their ability to produce IL-2, this could compromise Treg homeostasis, weaken peripheral tolerance and promote autoimmunity. Whether reduced IL-2 production as an early and consistent feature of T-cell responses in genetically at-risk individuals, and whether it precedes or predicts disease onset, remain an important question for future research. These findings contrast with the robust IL-2 production observed in treated CeD patients following oral gluten challenge, where circulating IL-2 levels can exceed 100 pg/ml within 2-4 hours of gluten ingestion and remain elevated for up to 6 hours (3, 39). In that setting, IL-2 is secreted by gluten-specific memory CD4^+^ T-cells and likely supports the expansion of these pathogenic effector cells. Given that treated CeD patients often experience small, unintentional gluten exposures (typically less than 1-2 g), it is plausible that this leads to intermittent, low-level IL-2 release. Such episodic IL-2 release could influence the broader T cell compartment, including naïve T cells. Whether contributes to altered homeostatic programming of naïve T cells over time warrants further investigation.

Finally, it is increasingly evident that CeD shares several immunogenetic features with other autoimmune conditions, including Type 1 diabetes, rheumatoid arthritis, multiple sclerosis and autoimmune thyroid disease. These disorders have overlapping risk loci that affect IL-2 signalling, T-cell activation and co-stimulation, and cytokine response pathways (40–42). The high prevalence of autoimmune comorbidities among individuals with CeD further supports the idea of shared underlying defects in immune regulation (13, 43). In this context, applying the T cell momentum assay to other autoimmune cohorts could provide valuable insight into whether dysregulation of naïve T-cells represents a generalisable early immune signature of autoimmunity.

Our findings suggest that intrinsic programming of naive CD4^+^ T cells is fundamentally altered in individuals with CeD. These cells exhibit defective integration of activation signals, impaired IL-2 secretion, and reduced proliferative momentum. The application of our momentum assay was instrumental in resolving these dynamic features and provides a valuable platform for dissecting early functional abnormalities in proliferative timers of division, death and DD in human T cells. Armed with this assay, our findings set the scene to explore whether naïve T-cell dysfunction is an important preclinical immune phenotype that predicts or contributes to the pathogenesis of CeD and other autoimmune disorders precursor in at-risk individuals.

Altogether, our findings support a model in which subtle, cell-intrinsic defects in signal integration and fate programming in naïve CD4^+^ T cells contribute to the initiation or amplification of pathogenic gluten-specific responses. The T cell momentum assay offers a new platform to characterise early immune abnormalities and may have broad utility in understanding T-cell dysfunction across a spectrum of autoimmune and immune-mediated diseases. Our results raise the intriguing possibility that altered programming of the naïve T cell compartment, manifesting as impaired IL-2 secretion and prolonged activation, may predispose individuals to the emergence of pathogenic CD4^+^ T-cell responses that are central to celiac disease.

## Methods

### Celiac and Healthy participants

Blood samples from individuals with CeD were sourced from the Celiac Research Database. A total of 22 CeD participants donated whole blood [median age, 44 years; range, 20-69 years; female, 19 (86%)]. Of these, 16 participants were following a strict gluten free diet for at least 6 months (treated CeD) and 6 donors were recently diagnosed with CeD (active CeD). All CeD participants had a confirmed diagnosis based on positive CeD serology and characteristic duodenal histopathology. Individual clinical details are described in Table 1. A total of 54 healthy donors provided blood samples (median age, 48 years; range, 21-75 years; female 23, 43%). Of these, 18 buffy coats were sourced from the Australian Red Cross LifeBlood (ARCLB), and 36 whole blood samples were sourced from the Volunteer Blood Donor Registry (VBDR), WEHI. Peripheral blood mononuclear cells (PBMCs) were isolated from all whole blood samples and used across all assays. Healthy donors were self-reported to be free of immune-mediated diseases, not taking immune-modulatory medication, and not acutely unwell.

**Table 1:** Clinical Information, HLA haplotype and demographics of Celiac Disease participants.

| Patient ID | Disease status | Age | Sex | Age Diagnosis | Other AI | Family Disease Hx | Medications | HLA DQ Haplotype (zygosity) | Patient PMHx |
| --- | --- | --- | --- | --- | --- | --- | --- | --- | --- |
| P01 | Treated | 64 | F | 49 | T1DM | Pernicious anaemia (mat. grandmother); Hashimoto's disease (mother) | Nil of note | DQ 2.5/8 | Nil of note |
| P02 | Treated | 61 | F | 51 | Nil Known | Graves' Disease (mother) | Nil of note | DQ 2.5 (Hom) | Nil of Note |
| P03 | Treated | 69 | F | 64 | Autoimmune thyroiditis | CeD (1 Brother, 3 nephews); Addison's disease (brother); Rheumatoid arthritis (sister) | Thyroxine/ Antihypertensive | DQ 2.5 (Hom) | Nil of note |
| P04 | Treated | 53 | M | 49 | Nil Known | CeD (mother, nephew) | Nil of note | DQ 2.2/2.5 | Abnormal LFTs |
| P05 | Treated | 66 | F | 62 | Graves' disease | CeD (daughter) | Nil of note | DQ 2.5 (Het) | Nil of note |
| P06 | Treated | 51 | F | 47 | Nil Known | Nil Known | Thyroxine 75 | DQ 2.5 Het | Participant (mother and sister) have "underactive thyroid"; eczema |
| P07 | Treated | 27 | F | 24 | Nil Known | T1DM (brother); CeD (2 sisters) | Nil of note | DQ 2.5/8 | Nil of Note |
| P08 | Treated | 43 | F | 33 | Nil Known | CeD (mother, son) | Nil of note | DQ 2.2/8 | Nil of note |
| P09 | Treated | 41 | F | 40 | Nil Known | Nil Known | Nil of note | DQ 2.5 Het | Nil of note |
| P10 | Treated | 32 | F | 23 | Nil Known | Nil Known | Nil of note | DQ 2.5/2.2 | Haemochromatosis |
| P11 | Treated | 56 | F | 55 | Hashimoto's disease | Hashimoto's disease (mother); CeD (cousin) | Thyroxine | DQ 2.5 Het | Nil of note |
| P12 | Treated | 42 | F | 41 | Nil Known | CeD (father);<br>Rheumatoid arthritis (father);<br>CeD (aunt, uncle) | Nil of note | DQ 2.5/8 | Asthma |
| P13 | Treated | 43 | F | 41 | Nil Known | T1DM (uncle) | Nil of note | DQ 2.5 Het | Factor 5 Leiden, Polyarthritis |
| P14 | Treated | 66 | F | 56 | Nil Known | CeD (cousin) | Nil of note | DQ 2.5 Het | Nil of note |
| P15 | Treated | 63 | M | 43 | Nil Known | CeD (brother, 2 sons, daughter) | Nil of Note | DQ 2.5 Het | Nil of note |
| P16 | Treated | 79 | F | 2 | Dermatitis<br>herpetiformis | CeD (2 sisters, 1 niece);<br>T1DM (3 sisters, 1 brother);<br>Hashimoto's disease (2 sisters, 1 niece);<br>Graves' disease (mother) | Anti-<br>hypertensives | DQ 2.5 Het | Hypertension, atrial fibrillation |
| P17 | Active | 57 | F | 57 | Nil Known | CeD (son) | Nil of note | DQ 2.5 Het | Lichen sclerosus |
| P18 | Active | 32 | F | 32 | Nil Known | Hypothyroidism (mother) | Nil of note | DQ 2.5 Het | Endometriosis, anxiety |
| P19 | Active | 28 | F | 28 | Nil Known | Inflammatory Bowel Disease (mother,<br>aunt) | Nil of note | DQ 2.5 Het | Nil of note |
| P20 | Active | 29 | F | 29 | Nil Known | Nil Known | Nil of note | DQ 2.5 Het | Nil of note |
| P21 | Active | 36 | F | 36 | Nil Known | CeD (brother) | Nil of note | DQ 2.5 Het | Asthma, rosacea, osteopaenia |
| P22 | Active | 47 | M | 47 | Nil Known | CeD (mat. aunt) | Nil of note | DQ 2.5/8 | Depression |

F, female; M, male; AI, autoimmunity; CeD, Celiac Disease; T1DM, Type 1 Diabetes Melitus; Hx, history; Mat, maternal; LFT, Liver function tests; Het, Heterozygous; Hom, Homozygous; PMHx, past Medical History

### Human Naïve CD4^+^ and CD8^+^ T lymphocyte culture

PBMCs were isolated using Ficoll-Paque Plus density gradient centrifugation in 50 ml Leucosep tubes (Greiner), washed and cryopreserved in FBS containing 10% (v/v) DMSO. For assay set up, PBMCs were thawed in a 37^°^C water bath and resuspended in warm T cell medium (TCM) (RPMI-1640 supplemented with 10% (v/v) FBS, 10 mmol/L HEPES, 100 U/ml Penicillin, 100 μg/ml streptomycin, 2 mmol/L GlutaMAX, 0.1 mmol/L non-essential amino acids, 1 mmol/L sodium pyruvate (Gibco, Thermo Fisher Scientific), 50 μM 2-mercaptoethanol, 0.1 mg/ml Normocin (InVivogen, CA, USA.)). Naive CD4^+^ or naive CD8^+^ T cells were isolated using EasySep Human Naïve CD4^+^ T Cell Isolation Kit II or EasySep Human CD8^+^ T Cell Isolation Kit (STEMCELL technologies) according to manufacturer’s instructions. Post isolation purities were ∼96% for naïve CD4^+^ T cells (CD3^+^CD20^-^CD4^+^CD8^-^CD45RA^+^CD45RO^-^CD27^+^) and ∼95% for naïve CD8^+^ T cells (CD3^+^CD20^-^CD4^-^ CD8^+^CD45RA^+^CD45RO^-^CD27^+^) as determined by flow cytometry using a monoclonal antibody panel (Supplemental Table 1; Supplementary Figure 1). Isolated naïve T cells were labelled with 5 μM division tracking dye CTV (Invitrogen) in PBS with 0.1% BSA for 20 min at 37^°^C. Cells were washed twice in TCM and kept on ice until stimulation.

CTV labelled naive CD4^+^ or CD8^+^ T cells were cultured in TCM and stimulated with anti-CD3/CD28 coated human activator Dynabeads (Gibco, Thermo Fisher Scientific) at a 1:1 cell-to-bead ratio with recombinant human IL-2 (100 U/ml). Cells were seeded into 96-well round-bottom plates (BD Falcon) at 1 x 10^4^ cells/well for CD4^+^ and 2ξ10^4^ cells/well for CD8^+^; and incubated for 42 hours at 37^°^C in 5% CO_2_. A subset of cells was harvested at 42 hours for flow cytometric analysis of kinetic parameters. Following activation, after 42 hours, Dynabeads were removed using magnetic separator and cells washed 3 times to remove residual rhuIL-2. Cells were then replated at 1 x 10^4^ cells/well and cultured for an additional four days under one of the following conditions: *(i)* media alone; *(ii)* anti-IL-2/IL-2Rα blockade: 5 μg/ml anti-human IL-2 antibody (clone MQ1-17H92, WEHI monoclonal antibody facility) plus 1 μg/ml anti-CD25 (Simulect Basiliximab; kindly provided by Novartis, Basel, Switzerland), *(iii)* IL-2 supplementation following blockade: anti-IL-2/anti-CD25 as above plus 31.6 U/ml rhuIL-2. Cells were harvested every 24 hours and analysed by flow cytometry. Additional aliquots were stained for activation markers (CD69, CD62L, CD127) and analysed using flow cytometry (Supplemental Table 2).

### Treg stain and quantification from PBMCs

Cryopreserved PBMCs were thawed in pre-warmed TCM, washed, and resuspended in PBS/10% FBS. Cells were filtered through a 70 μm cell strainer to remove dead cells, then resuspended at 2ξ10^7^ cells/ml. Fc Block (BD Biosciences) was added according to manufacturer’s instructions and incubated for 10 min at room temperature. Cells were aliquoted into v-bottom 96-well plates, centrifuged and supernatant carefully removed. PBMCs were then stained with a pre-mixed Treg antibody cocktail containing (Supplemental Table 3), on ice in the dark for 30 min, washed twice and analysed on a BD Fortessa X20 flow cytometer. Tregs (CD3^+^CD4^+^CD8^-^CD25^+^CD127^-^) were determined as the percentage of lymphocytes within the live lymphocyte gate.

### IL-2 secretion assay and CD25 surface expression quantification

Naïve CD4^+^ and CD8^+^ T cells were stimulated with anti-CD3/CD28 coated human activator Dynabeads in the absence of rhuIL-2 then supernatant was collected at the 42-hour timepoint. In a parallel condition, cells were stimulated with anti-CD3/anti-CD28 with 100U/mL rhuIL-2 for 42 hours, followed by bead and supernatant removal, then re-cultured in media alone for a further 24 hours (70 hours total), after which supernatant was again collected. Remaining cells from both conditions were stained with anti-human CD25-BV421 (Biolegend) and gMFI of CD25 cell surface expression was quantified using flow cytometry.

IL-2 concentration in cell culture supernatants was measured using V-plex Human IL-2 (Meso Scale Diagnostics) according to manufacturer’s instructions. Detection was performed on a MESO QuickPlex SQ 120 plate reader using Discovery Workbench 4.0 software. The lower limit of detection (LLOD) was calculated for each plate and was approximately 0.1 pg/ml.

### Flow Cytometry

Proliferation kinetics were analysed using CTV-labelled T cells on a BD Canto II (Becton Dickinson). To enable absolute cell quantification, CaliBRITE rainbow calibration particles (BD Biosciences) were added at 1×10^4^ beads per well immediately prior to acquisition. Dead cells were excluded using propidium iodide (0.2 μg/ml, Sigma-Aldrich).

For T-cell activation markers, IL-2Rα (CD25) expression and Treg quantification, samples were analysed on a BD LSRFortessa X20 cytometer. Hydroxystilbamidine methanesulfonate (1μg/ml; Invitrogen, Thermo Fisher Scientific) was added immediately prior to acquisition for dead cell discrimination. All flow cytometry data were analysed using FlowJo software (Treestar) including gating strategies, population statistics and gMFI calculations.

### Cell counting and Precursor cohort method

Total cell numbers were determined by multiplying the number of events acquired with the ratio of total rainbow calibration particles added and amount collected by the flow cytometer:

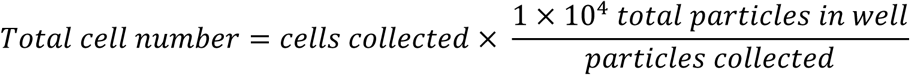

Cell survival and division were quantified using the previously established precursor cohort method (25). The precursor cohort or initial pre-stimulated undivided cells that contributed to each CTV peak were calculated as:

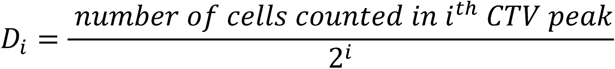

Where *i* is the division generation number.

The total Cohort number and Mean Division Number (MDN) were calculated as:

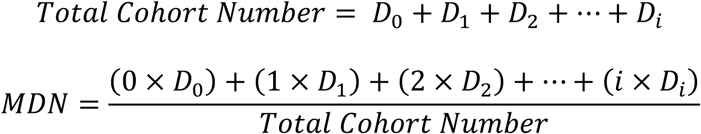

Total cell numbers and total cohort numbers were expressed as fold-change relative to initial cell seeding conditions; day 0 for 42-hour values, or day 2 for time points after stimulus removal. The Area Under the Curve (AUC) was calculated using the normalised total cell number (relative to day 2) over the course of the experiment (day 2 to day 6). AUC was estimated using the standard trapezoidal rule for non-uniformly spaced data points, {(*t*_0_, *c*_0_), (*t*_1_, *c*_1_), …, (*t*_*n*_, *c*_*n*_)}, where *t*_*j*_ and *c_j_* are *j*^th^ harvested time point and normalised total cell number, respectively, such that AUC = 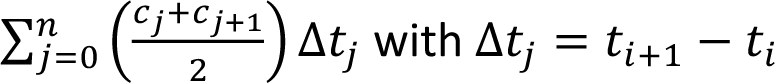

### Cyton2 Modelling

To model T cell population dynamics, we applied the Cyton2 framework and a standard fitting strategy to accurately capture the underlying cellular division timers (16). The model incorporates three lognormally distributed random variables (RVs): first division 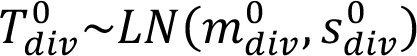, time to death 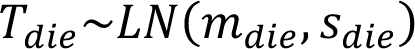 and time to division destiny 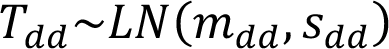 where (*m*, *s*) denote median and log-variance, respectively, of the lognormal distribution. A constant *b* ∈ ℝ_>0_ represents the average time between subsequent divisions. The model was fitted to observed cell counts, denoted 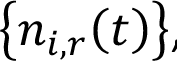, where: 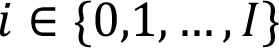 represents generation, 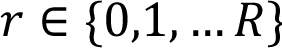 is a replicate index, 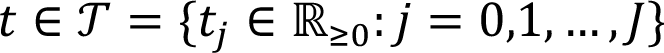 is discrete time points, and 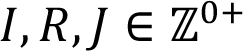 are the maximum observed generation, total number of replicates and final time point, respectively. The model contains 7 parameters: 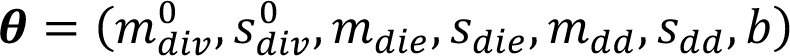 for each set of measurements. We define the following cost function to minimise the residual sum-of-squares (RSS),

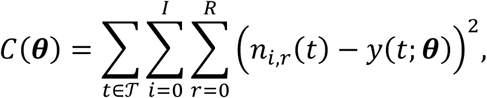

where *y*(*t*; **θ**) is the model prediction. The minimisation and best-fit parameters were achieved by utilising the least-squares method and Levenberg-Marquardt optimisation algorithm (44) such that

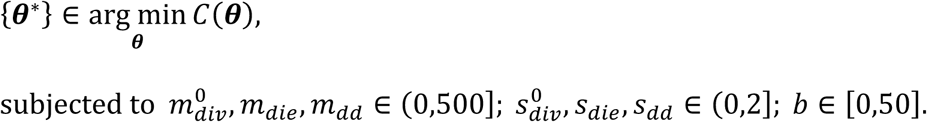

The best-fit parameters were selected based on the minimum RSS. This process requires a set of parameter guesses to initiate the algorithm. Consequently, 100 uniform random sets of initial guesses were assigned for each parameter component within the identified ranges. Then, the best-fit parameters were determined based on the corresponding lowest RSS value. Bootstrap resampling (n=1000) was used to estimate 95% confidence intervals for the fitted parameters and generate confidence bands for model-predicted cell numbers (45). Specifically, we constructed 1000 artificial datasets by resampling with replacement from the original dataset at each time point, allowing us to obtain an additional 1000 estimates to compute the errors.

## Statistical Analysis

Due to the use of cryopreserved PBMCs across multiple experiments, not all donors were represented in each dataset. For active CeD samples, only CD4^+^ T cell results were generated due to sample limitations. All statistical analyses were performed using GraphPad Prism v9.5.0 unless otherwise stated. Welch’s t-test was applied for parametric comparisons. Linear and non-linear curve-fitting were applied where appropriate. Correlation analyses used Spearman’s rank method. Statistical significance was defined as *p* < 0.05. For Cyton2 model parameters, a custom non-parametric permutation test was implemented using Python v3.11.5 to evaluate differences between healthy and CeD donor estimates. Each parameter was assessed independently with null and alternative hypotheses defined as: 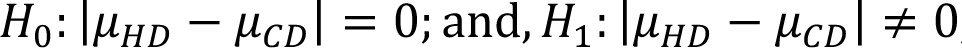, where μ_*HD*_ and μ_*CD*_ are the mean parameter estimates for healthy and CeD donor samples, respectively. Two-sided *p-*values were calculated using Monte-Carlo simulations with 1 x 10^?^ random permutations.

## Study approvals

Ethical approval for all study protocols was granted by the Human Research Ethics Committees of Melbourne Health HREC/16/MH261, 2009.162, and 2020.162 and WEHI (10/02). All participants provided written, informed consent in accordance with the Declaration of Helsinki and its later revisions.

## Data Availability

Supporting data values for each figure and supplemental figure are provided in the Supporting Data Values spreadsheet. Precursor cohort data files, including individual donor samples, cell types (CD4^+^ or CD8^+^) and conditions (media alone, anti-IL-2/IL-2Rα and anti-IL-2/IL-2Rα + rhuIL-2), used to generate proliferation analyses and Cyton2 modelling outputs, are available in the following GitHub Repository (https://github.com/hodgkinlab/CeDmomentum2025). This repository also includes Cyton2 model parameter estimates (medians and standard errors) for each cellular timer and figures of the fitted lognormal distributions for all donor responses.

## Author Contributions

Conceptualisation: V.L.B, P.D.H and S.H

Methodology: A.J.F, H.C, S.H, G.M.V, J.N, M.M, D.V, M.Y.H, P.D.H and V.L.B

Conducting experiments & data acquisition: A.J.F, H.C, G.M.V, J.N, M.M, D.V, L.J.H, M.Y.H, L.M.H, M.F

Data analysis and software: A.J.F, H.C, S.H, P.D.H and V.L.B

Visualisation: A.J.F, H.C

Original draft: A.J.F, H.C, S.H, J.A.T-D, P.D.H and V.L.B

Funding Acquisition: V.L.B, P.D.H and J.A.T-D

Supervision: V.L.B, P.D.H, S.H

## Acknowledgements

We thank all study participants from VBDR, ARCLB and those with celiac disease for their willingness and time to donate blood samples. We thank staff at ARCLB for acquiring blood from healthy donors and assisting with donor details. We thank Amy Russell and Olivia Moscatelli from the Tye-Din laboratory for their support in running MSD IL-2 assays. We thank Novartis for the kind gift of anti-CD25 mAb used in this study. We gratefully acknowledge and appreciate the support and services provided by the WEHI Clinical Discovery and Translation, WEHI Antibody Facility, and WEHI Flow Cytometry unit.

