## Supplemental Figures 1-9; Supplemental table 1-3 for "Functional immune profiling reveals CD4^+^ T cell dysregulation associated with coeliac disease"

#### Supplemental Figure Legends

**Supplemental Figure 1. Gating strategies for naïve T cells purity assessment and CTV division profiles.**

**(A)** Human PBMCs and isolated naïve T cells were stained with lineage and viability markers to assess purity of naïve CD4$^+$ T cells by flow cytometry. Naïve CD4+ T cells were defined as CD3$^+$CD20$^-$CD4$^+$CD8$^-$CD45RA$^+$CD45RO$^-$CD27$^+$ and naïve CD8$^+$ T cells as CD3$^+$CD20$^-$CD4$^-$CD8$^+$CD45RA$^+$CD45RO$^-$CD27$^+$. Total PBMCs (green overlay) were used to set gates appropriately. The gating strategy proceeded sequentially through lymphocytes, single cells, live cells, CD3$^+$CD20$^-$ T cells, then CD4$^+$ (red) or CD8$^+$ (blue) subsets, and finally CD45RA$^+$CD45RO$^-$CD27$^+$ to identify naïve cells. **(B)** For each timepoint, counting particles and PI were added to cultures to quantify CTV dilution and determine the percentage of unstimulated cells. Gating strategy involved gating in the order of lymphocytes, single cells, live cells, followed by CTV histograms to determine division generation (D0-D8+), or gating small, undivided, and unstimulated cells as shown.

**Supplemental Figure 2. Optimisation of Dynabead stimulation, cell density, IL-2 blockade, and rhuIL-2 supplementation in the naïve T cell momentum assay.**

**(A)** Naïve CD4$^+$ T cells ($1x10^4$) were stimulated with anti-CD3/CD28 Dynabeads at 1:2, 1:1, or 2:1 bead-to-cell ratios. Total cell number (left), total cohort number (middle) and mean division number (MDN; right) were quantified over 72 hours. **(B)** Varying input densities of 5,000, 10,000 and 15,000 naïve CD4$^+$ T cells were cultured with 1:1 bead ratio. Fold change in total cell numbers, total cohort number and MDN were quantified over 72 hours. **(C)** Naïve CD4$^+$ T cells were stimulated for 42 hours using a 1:1 bead ratio and 100 U/ml rhuIL-2, stimuli were removed and cells recultured in anti-IL-2/IL-2R$\alpha$ blocking antibodies at varying cell densities (5,000-15,000 cells/well).

Expansion and division metrics were tracked until ~144 hours. **(D)** Activated naïve $CD4^+$ T cells were cultured with 10 or 100 U/ml rhuIL-2 in the presence of titrated anti-IL-2 or anti-CD25 for 24 hours. Metabolic activity was assessed alamarBlue fluorescence (excitation/emission: 550/590 nm) to determine optimal blocking concentrations (highlighted in red). **(E)** Following initial activation, naïve $CD4^+$ T cells were cultured in the momentum assay in anti-IL-2/IL-2R$\alpha$ with titrated rhuIL-2 concentrations (0-316 U/ml) Fold change in total cell number, total cohort number and MDN were assessed over time. **(F-G)** Overlayed, modally normalised CTV profiles from day 4 (black) and day 6 (purple) are shown for $CD4^+$ T cells cultured in either (F) anti-IL-2/IL-2R$\alpha$ alone or (G) anti-IL-2/IL-2R$\alpha$ + rhuIL-2. **(H)** Change in MDN ($\Delta$ MDN) from day 4 to day 6 in these two conditions was calculated to assess IL-2 dependent proliferation. **(I)** Myc protein expression (gMFI) was measured in activated $CD8^+$ T cells stimulated using the momentum assay then recultured in media alone, anti-IL-2/IL-2R$\alpha$, anti-IL-2/IL-2R$\alpha$ with rhuIL-2, or rhuIL-2 alone, as indicated. Data are presented as means of fold change total cell numbers, total cohort numbers, MDN, fluorescence (FU) and myc gMFI $\pm$ SD of duplicate cultures shown. Data is representative of 2-3 independent experiments. Means of $\Delta$ MDN $\pm$ SEM of anti-IL-2/IL-2R$\alpha$ and anti-IL-2/IL-2R$\alpha$ + rhuIL-2 conditions shown.

**Supplemental Figure 3. Sex comparisons of $CD4^+$ and $CD8^+$ T-cell responses**

Human naïve T cells from HDs were stimulated using the momentum assay, and responses were assessed at 42-hours and after stimulus removal. **(A)** Sex comparisons in $CD4^+$ T-cell response, including cell survival, division, and activation, at 42 hours (top panels) and after stimuli removal (bottom panels); n= 13 males, 9 females). **(B)** Equivalent analysis of $CD8^+$ T-cell responses in male and female donors at 42 hours (top panels) and after stimuli removal (bottom panels); n = 20 males, 12 females). Data shown include mean $\pm$ SEM for fold change in total cell number, total cohort number, MDN, percentage of unstimulated cohort, and AUC. Statistical comparisons were performed using unpaired t-test with Welch's correction.

**Supplemental Figure 4. Age-related analysis of naïve T-cell responses**

Correlations between donor age and T-cell responses were assessed in HD. For $CD4^+$ T cells: **(A)** AUC, (left) Fold change in total cohort number (day 6, middle), and (right) maxMDN. **(B)** For $CD8^+$ T cells: AUC (left), Fold change total cohort number (day 6, middle), and maxMDN (right panel). **(C)** Violin plots showing the age distribution (median and quartiles) of HD and CeD donors included in the $CD4^+$ (HD = 22, CeD = 17), and $CD8^+$ datasets (HD = 32, CeD = 15).

**Supplemental Figure 5. Stratification of CeD donor responses by disease activity and analysis of unstimulated naïve T cell survival**

**(A)** Naïve $CD4^+$ T cells from healthy, gluten free (GF) and active (Act) CeD donors were stimulated with anti-CD3/CD28 beads and rhuIL-2 for 42 hours. Proliferative responses were assessed by activation, response, survival and division, as measured by percent of unstimulated cohort, fold change in total cell number, total cohort number, and MDN, respectively (N: HD = 22, GF = 12, Act = 5). **(B)** After initial stimulation, $CD4^+$ T cell proliferation momentum was tracked over an additional 4 days in the same donor groups. Fold change in total cell number, total cohort number and MDN were quantified to assess proliferation dynamics. **(C-D)** Naïve $CD4^+$ and $CD8^+$ T cells from HD and CeD donors were cultured in media without stimuli. Cell numbers were assessed over time by flow cytometry to calculate fold change and half-life ($t^{1/2}$) of individual donor survival curves ($CD4^+$: HD = 6, CeD = 5; $CD8^+$: HD = 8, CeD = 6). Data are presented as means $\pm$ SEM for percent unstimulated, fold change in total cell number, total cohort number, MDN and survival $t^{1/2}$. Comparisons were made using unpaired t-tests with Welch's correction. * indicates $p<0.05$; ** indicates $p<0.01$.

**Supplemental Figure 6. Comparable CD8$^+$ T cell response momentum in HD and CeD donors**

Naïve CD8$^+$ T cells from HD and CeD individuals were stimulated for 42 hours as above, stimuli removed, and cells were recultured in different conditions. Cells were harvested and analysed by flow cytometry every 24 hours until day 6. **(A-C)** Cells recultured in media alone **(A),** with anti-IL-2/IL2R$\alpha$ **(B)**, or with anti-IL-2/IL-2R$\alpha$ and 31.6 U/mL rhuIL-2 **(C)**. For each condition, fold change total cell numbers (left panels), total cohort numbers (middle panels), and MDN (right panels) for CD8$^+$ T cells is shown. **(D-F)** Summary metrics comparing media, anti-IL-2/IL-2R$\alpha$, and anti-IL-2/IL-2R$\alpha$ + rhuIL-2 conditions: AUC for total cell number (D), day 6-fold change in survival (E) and division (F). Data are shown as mean $\pm$ SEM for HD and CeD groups shown (N: HD = 33-35, CeD = 15-16). Comparisons used unpaired t-tests with Welch's correction. * indicates p<0.05.

**Supplemental Figure 7. Cyton2 modelling reveals no significant differences in cellular division dynamics between HD and CeD under IL-2 blockade.**

Cyton2 modelling was used to evaluate T cell division kinetics from HD and CeD donors when IL-2 signalling was blocked. **(A)** Empirical cumulative distribution functions (eCDFs) for estimated medians timers and subsequent division times of CD4+ T cells under IL-2 blockade. Each step-up represents an individual donor (HD: green, CeD: red). **(B)** Permutation test results and null distributions for CD4$^+$ T cells. Histogram show test statistics from $10^7$ random permutations; vertical dashed line indicates the observed statistic, with two-sided $p$ value shown. **(C, D)** eCDFs (C) and permutation test (D) analyses for CD8$^+$ T cells. **(E)** Timer medians and subsequent division times were compared between HD and CeD cohorts for CD4$^+$ and CD8$^+$ T cell responses. Data shown are mean $\pm$ SEM of HD and CeD groups (CD4$^+$ N: HD = 22, CeD = 16; CD8$^+$ N: HD = 3,4 CeD = 15). Comparisons used unpaired t-tests with Welch's correction.

**Supplemental Figure 8. IL-2 secretion and CD25 expression in activated naïve $CD8^+$ T cells and CeD IL-2 subgroup analysis.**

Supernatants and cells were collected from $CD4^+$ and $CD8^+$ T cell cultures stimulated using the momentum assay at Day 3 or from cultures containing anti-CD3/CD28 beads alone at 42 hours. (A) IL-2 concentrations were measured in supernatants and (B) cells were stained to measure surface expression of high affinity IL-2R$\alpha$ (CD25) by flow cytometry at day 3 of the momentum assay (*n*: HD = 11, CeD = 7). **(C)** IL-2 concentration (N: HD = 11, CeD = 8) and **(D)** CD25 expression (N: HD = 6, CeD = 4) were quantified at 42 hours when $CD8^+$ T cells where cultured in anti-CD3/CD28 beads alone. **(E)** IL-2 secretion was assessed in cultures containing $CD4^+$ T cells from healthy, GF and active CeD at day 3 of the momentum assay (N: HD = 23, GF = 9, Act = 5) and **(F)** 42 h with anti-CD3/CD28 beads alone (N: HD = 19, GF = 9, Act = 5). Data shown as mean $\pm$ SEM for IL-2 concentration and IL-2R$\alpha$ (CD25) gMFI. Comparisons used unpaired t-tests with Welch's correction. * indicates $p<0.05$

**Supplemental Figure 9. Flow cytometry gating strategies and T cell activation marker analysis**

**(A)** Treg gating strategy: PBMCs were stained with Fluorogold and antibodies to identify T regulatory cells ($CD3^+CD4^+CD8^-CD25^{hi}CD127^{lo}$). Gating strategy involved gating in the order of lymphocytes, single cells, live cells, $CD3^+$ T cells, $CD4^+CD8^-$ cells, $CD25^{hi}CD127^{lo}$ cells. **(B)** T-cell activation marker gating: naïve $CD4^+$ T cells stimulated via the T cell momentum assay were stained for CD69, CD25, CD62L and CD127, followed by live/dead dye (Fluorogold). Gating strategy involved gating in the order of lymphocytes, single cells, live cells, then CTV vs each activation marker (CD69, CD25, CD62L or CD127). **(C)** gMFI of CD62L (left) and CD127 (right) was quantified across the response (N: HD = 8-9, CeD = 9). **(D)** %$CD69^+$ cells on day 3 of the of the momentum assay for $CD8^+$ T cells (left; N: HD = 11, CeD = 7) and $CD4^+$ T cells (right) was determined for healthy, GF and active

CeD individuals (N: HD = 16, GF = 9, Act = 5). Data presented as mean ± SEM. Comparisons used unpaired *t* tests with Welch's correction. * indicates $p<0.05$; **** indicates $p<0.0001$

*Supplemental Table 1: Human Naïve T cell purity monoclonal antibody panel.*

| Antibody (anti-human) | Fluorochrome | Clone | Supplier | Catalogue Number |
|---|---|---|---|---|
| CD20 | BUV395 | 2H7 | BD Biosciences | 563782 |
| CD3 | PacBlue | UCHT1 | BD Biosciences | 558117 |
| CD4 | APC | RPA-T4 | BD Biosciences | 555349 |
| CD8 | APC-780 | SK1 | eBioscience | 47-0087-42 |
| CD45RA | PE-Cy7 | HI100 | eBioscience | 25-0458-42 |
| CD45RO | PE | UCHL1 | eBioscience | 12-0457-42 |
| CD27 | FITC | MT271 | Miltenyi Biotec | 130-113-634 |

*Supplemental Table 2: Human T cell activation marker panel.*

| Antibody (anti-human) | Fluorochrome | Clone | Supplier | Catalogue Number |
|---|---|---|---|---|
| CD69 | FITC | FN50 | Biolegend | 310904 |
| CD62L | PE-Cy7 | DREG-56 | Invitrogen | 25062942 |
| CD127 | BV650 | A019D5 | Biolegend | 351326 |

*Supplemental Table 3: Human T regulatory cell marker panel.*

| Antibody (anti-human) | Fluorochrome | Clone | Supplier | Catalogue Number |
|---|---|---|---|---|
| CD3 | BUV395 | SK7 | BD Biosciences | 564001 |
| CD25 | BV421 | BC96 | Biolegend | 302630 |
| CD127 | BV650 | A019D5 | Biolegend | 351326 |
| CD4 | PerCP-Cy5.5 | SK3 | BD Biosciences | 341654 |
| CD8 | APC-eF780 | SK1 | Invitrogen | 47008742 |

### Supplementary Figure 1

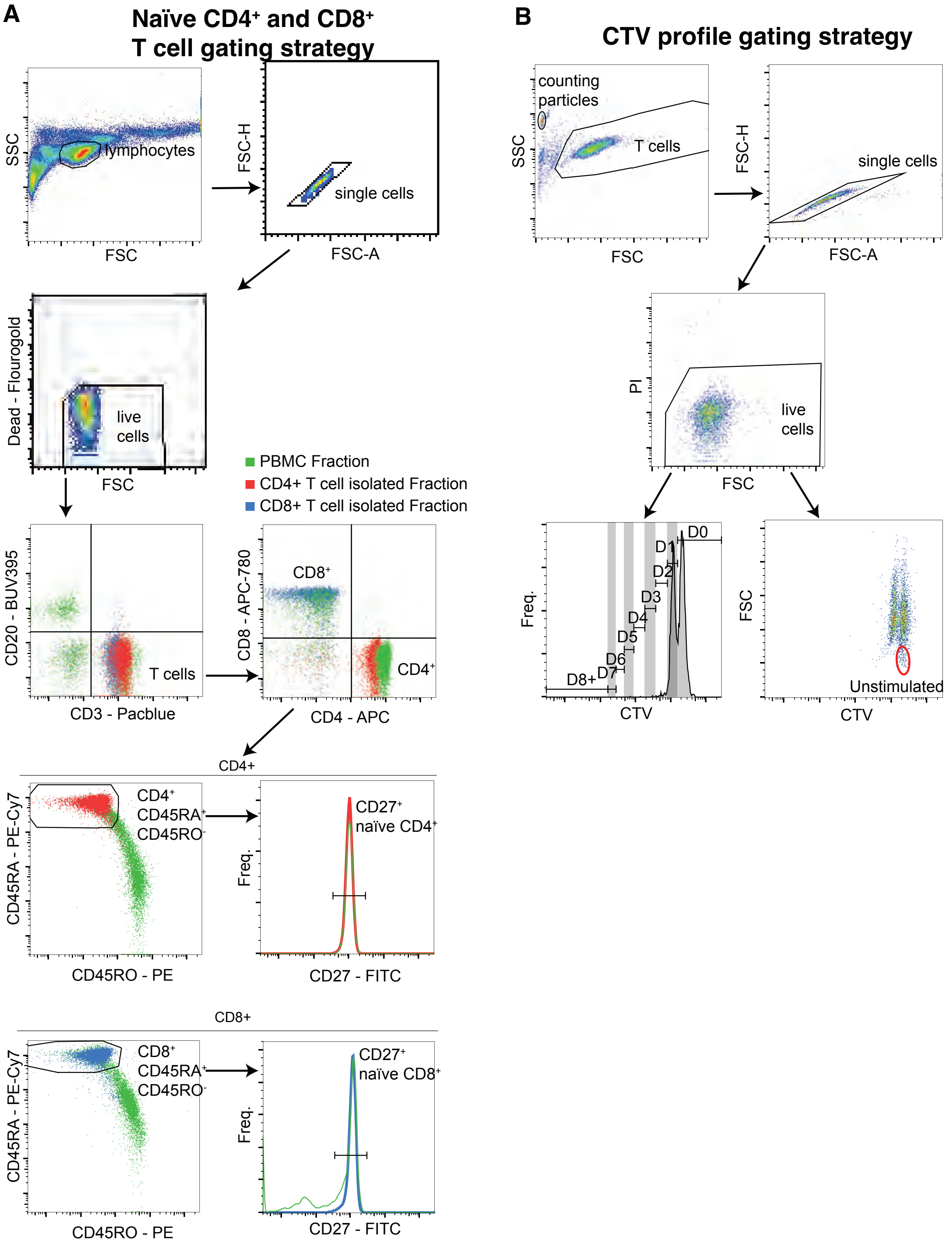


**Supplemental Figure 1. Gating strategies for naïve T cells purity assessment and CTV division profiles.** (A) Human PBMCs and isolated naïve T cells were stained with lineage and viability markers to assess purity of naïve $CD4^+$ T cells by flow cytometry. Naïve CD4+ T cells were defined as $CD3^+CD20^-CD4^+CD8^-CD45RA^+CD45RO^-CD27^+$ and naïve $CD8^+$ T cells as $CD3^+CD20^-CD4^-CD8^+CD45RA^+CD45RO^-CD27^+$. Total PBMCs (green overlay) were used to set gates appropriately. The gating strategy proceeded sequentially through lymphocytes, single cells, live cells, $CD3^+CD20^-$ T cells, then $CD4^+$ (red) or $CD8^+$ (blue) subsets, and finally $CD45RA^+CD45RO^-CD27^+$ to identify naïve cells. (B) For each timepoint, counting particles and PI were added to cultures to quantify CTV dilution and determine the percentage of unstimulated cells. Gating strategy involved gating in the order of lymphocytes, single cells, live cells, followed by CTV histograms to determine division generation (D0-D8+), or gating small, undivided, and unstimulated cells as shown.

### Supplementary Figure 2

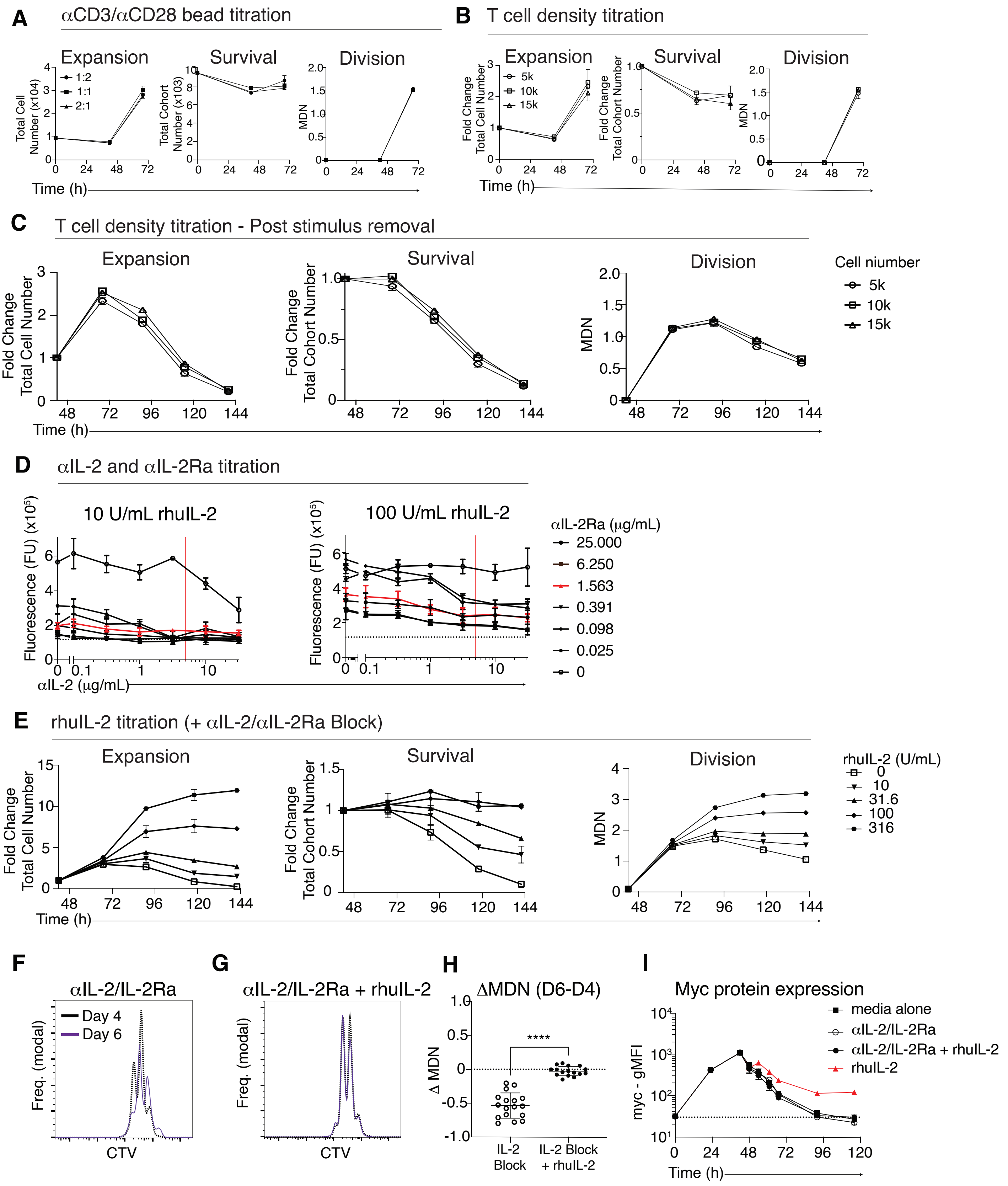

### Supplementary Figure 2

**Supplemental Figure 2. Optimisation of Dynabead stimulation, cell density, IL-2 blockade, and rhuIL-2 supplementation in the naïve T cell momentum assay.**
(A) Naïve CD4+ T cells (1x$10^4$) were stimulated with anti-CD3/CD28 Dynabeads at 1:2, 1:1, or 2:1 bead-to-cell ratios. Total cell number (left), total cohort number (middle) and mean division number (MDN; right) were quantified over 72 hours. (B) Varying input densities of 5,000, 10,000 and 15,000 naïve CD4+ T cells were cultured with 1:1 bead ratio. Fold change in total cell numbers, total cohort number and MDN were quantified over 72 hours. (C) Naïve $CD4^+$ T cells were stimulated for 42 hours using a 1:1 bead ratio and 100 U/ml rhuIL-2, stimuli were removed and cells recultured in anti-IL-2/IL-2Rα blocking antibodies at varying cell densities (5,000-15,000 cells/well). Expansion and division metrics were tracked until ~144 hours. (D) Activated naïve CD4+ T cells were cultured with 10 or 100 U/ml rhuIL-2 in the presence of titrated anti-IL-2 or anti-CD25 for 24 hours. Metabolic activity was assessed alamarBlue fluorescence (excitation/emission: 550/590 nm) to determine optimal blocking concentrations (highlighted in red). (E) Following initial activation, naïve $CD4^+$ T cells were cultured in the momentum assay in anti-IL-2/IL-2Rα with titrated rhuIL-2 concentrations (0-316 U/ml) Fold change in total cell number, total cohort number and MDN were assessed over time. (F-G) Overlayed, modally normalised CTV profiles from day 4 (black) and day 6 (purple) are shown for CD4+ T cells cultured in either (F) anti-IL-2/IL-2Rα alone or (G) anti-IL-2/IL-2Rα + rhuIL-2. (H) Change in MDN (ΔMDN) from day 4 to day 6 in these two conditions was calculated to assess IL-2 dependent proliferation. (I) Myc protein expression (gMFI) was measured in activated CD8+ T cells stimulated using the momentum assay then recultured in media alone, anti-IL-2/IL-2Rα, anti-IL-2/IL-2Rα with rhuIL-2, or rhuIL-2 alone, as indicated. Data are presented as means of fold change total cell numbers, total cohort numbers, MDN, fluorescence (FU) and myc gMFI ± SD of duplicate cultures shown. Data is representative of 2-3 independent experiments. Means of ΔMDN ± SEM of anti-IL-2/IL-2Rα and anti-IL-2/IL-2Rα + rhuIL-2 conditions shown.

### Supplementary Figure 3

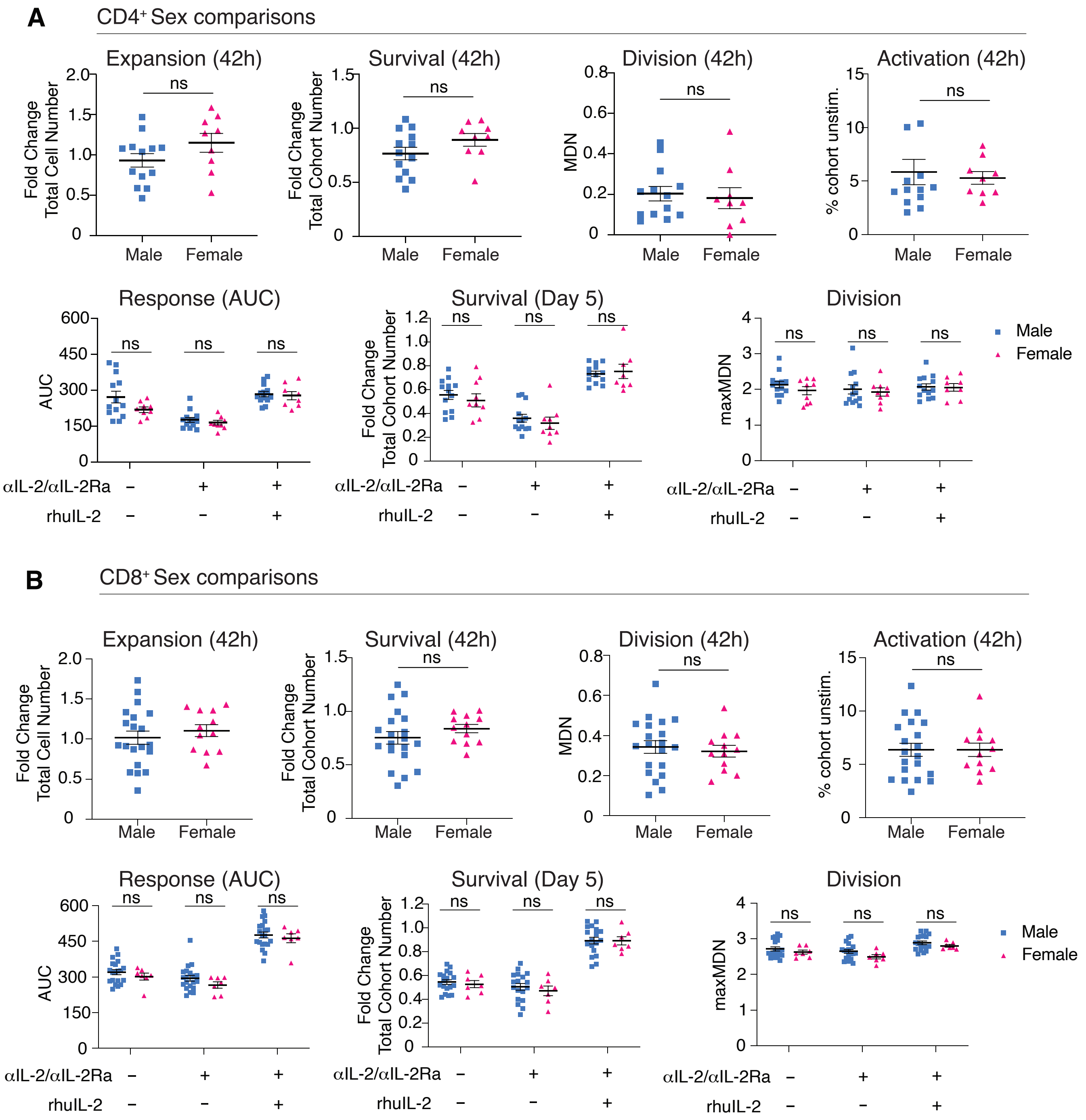


**Supplemental Figure 3. Sex comparisons of CD4⁺ and CD8⁺ T-cell responses**

Human naïve T cells from HDs were stimulated using the momentum assay, and responses were assessed at 42-hours and after stimulus removal. (A) Sex comparisons in CD4+ T-cell response, including cell survival, division, and activation, at 42 hours (top panels) and after stimuli removal (bottom panels); n= 13 males, 9 females). (B) Equivalent analysis of CD8⁺ T-cell responses in male and female donors at 42 hours (top panels) and after stimuli removal (bottom panels); n = 20 males, 12 females). Data shown include mean ± SEM for fold change in total cell number, total cohort number, MDN, percentage of unstimulated cohort, and AUC. Statistical comparisons were performed using unpaired t-test with Welch's correction.

### Supplementary Figure 4

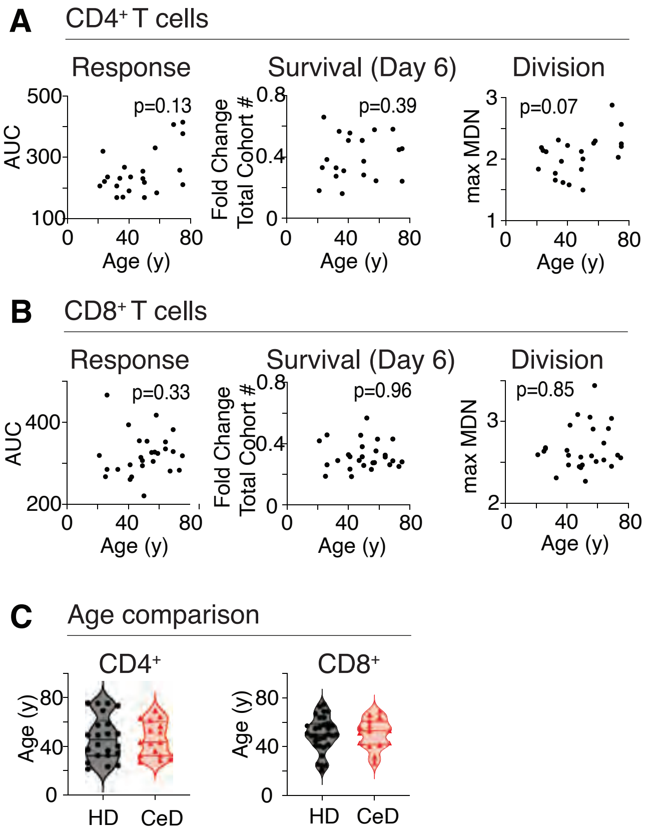


**Supplemental Figure 4. Age-related analysis of naïve T-cell responses**

Correlations between donor age and T cell responses were assessed in HD. For CD4+ T cells: **(A)** AUC, (left) Fold change in total cohort number (day 6, middle), and (right) maxMDN. **(B)** For CD8+ T cells: AUC (left), Fold change total cohort number (day 6, middle), and maxMDN (right panel). **(C)** Violin plots showing the age distribution (median and quartiles) of HD and CeD donors included in the CD4+ (HD = 22, CeD = 17), and CD8+ datasets (HD = 32, CeD = 15).

### Supplementary Figure 5

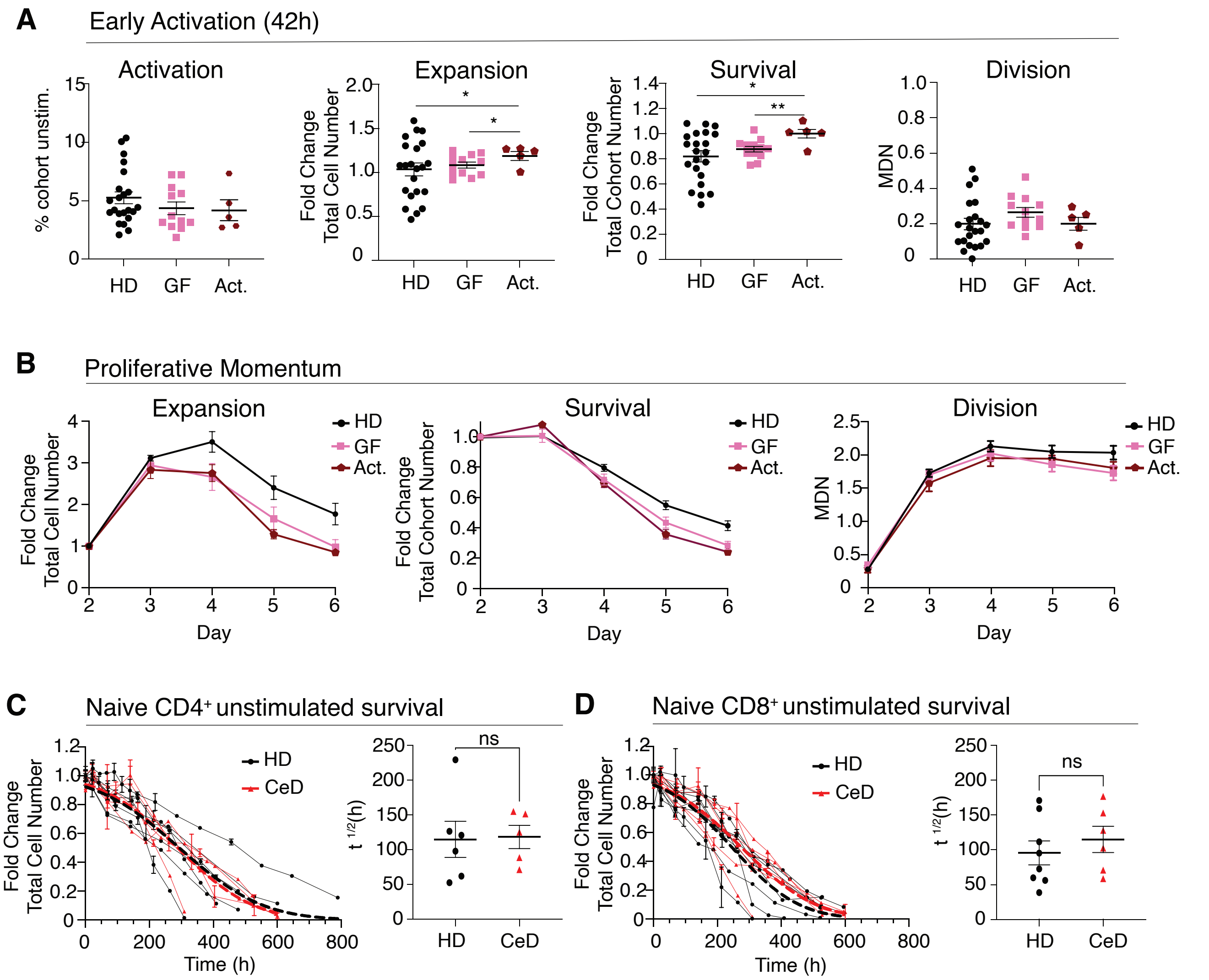


**Supplemental Figure 5. Stratification of CeD donor responses by disease activity and analysis of unstimulated naïve T cell survival**

**(A)** Naïve CD4+ T cells from healthy, gluten free (GF) and active (Act) CeD donors were stimulated with anti-CD3/CD28 beads and rhuIL-2 for 42 h. Proliferative responses were assessed by activation, response, survival and division, as measured by percent of unstimulated cohort, fold change in total cell number, total cohort number, and MDN, respectively (N: HD = 22, GF = 12, Act = 5). **(B)** After initial stimulation, $CD4^{+}$ T cell proliferation momentum was tracked over 6 days in the same donor groups. Fold change in total cell number, total cohort number and MDN were quantified to assess proliferation dynamics. **(C-D)** Naïve $CD4^{+}$ and $CD8^{+}$ T cells from HD and CeD donors were cultured in media without stimuli. Cell numbers were assessed over time by flow cytometry to calculate fold change and half-life ($t^{1/2}$) of individual donor survival curves ($CD4^{+}$: HD = 6, CeD = 5; CD8+: HD = 8, CeD = 6). Data are presented as means ±SEM for percent unstimulated, fold change in total cell number, total cohort number, MDN and survival $t^{1/2}$. Comparisons were made using unpaired t-tests with Welch's correction. * indicates p<0.05; ** indicates p<0.01.

### Supplementary Figure 6

#### CD8⁺ Momentum Assay Results

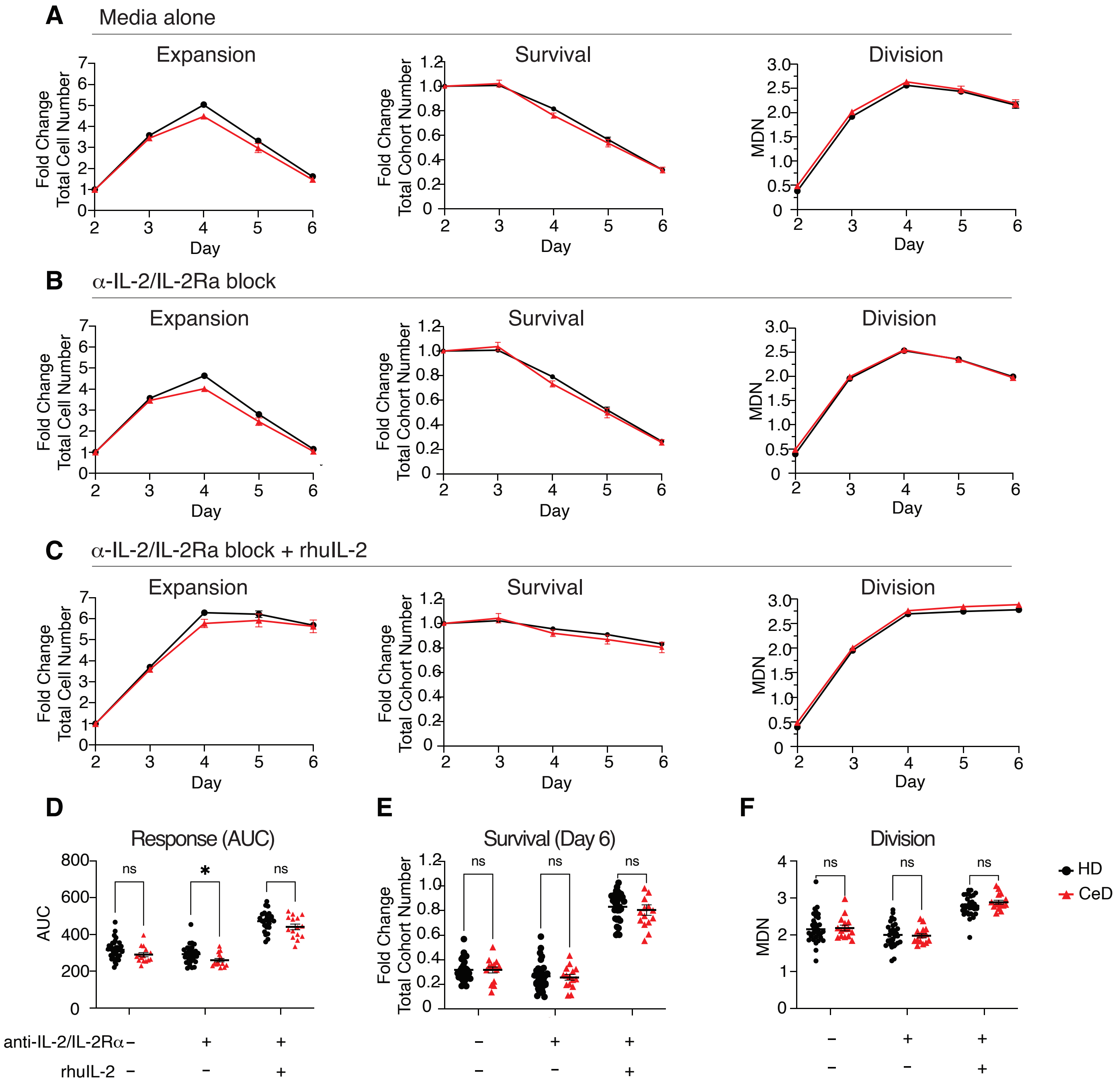


**Supplemental Figure 6. Comparable CD8+ T-cell response momentum in HD and CeD donors**

Naïve CD8+ T cells from HD and CeD individuals were stimulated for 42 hours as above, stimuli removed, and cells were recultured in different conditions. Cells were harvested and analysed by flow cytometry every 24 hours until day 6. **(A-C)** Cells were recultured in media alone **(A)**, with anti-IL-2/CD25 blocking antibodies **(B)**, **or** with anti-IL-2/IL-2Rα and 31.6 U/ml rhuIL-2 **(C)**. For each condition, fold change total cell numbers (left panels), total cohort numbers (middle panels), and MDN (right panels) for CD8+ T cells is shown. **(D-F)** Summary metrics comparing media, anti-IL-2/IL-2Rα and anti-IL-2/IL-2Rα + rhuIL-2 conditions: AUC for total cell number (D), day 6-fold change in survival (E) and division (F). Data are shown as mean ±SEM for HD and CeD groups shown (N: HD = 33-35, CeD = 15-16). Comparisons used unpaired t-tests with Welch's correction. * indicates $p<0.05$.

### Supplementary Figure 7

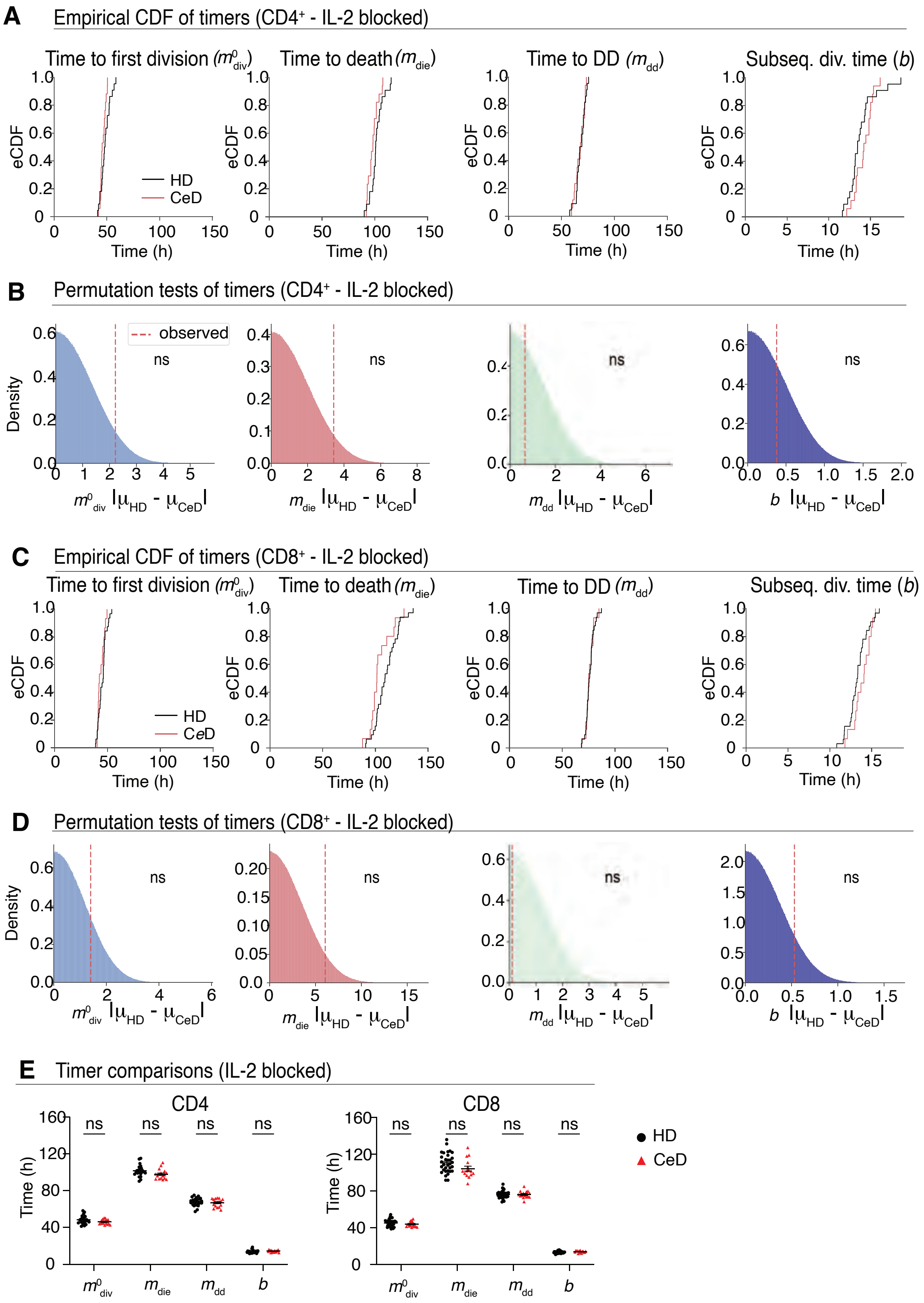

### Supplementary Figure 7

**Supplemental Figure 7. Cyton2 modelling reveals no significant differences in cellular division dynamics between HD and CeD under IL-2 blockade.**

Cyton2 modelling was used to evaluate T cell division kinetics from HD and CeD donors when IL-2 signalling was blocked. (A) Empirical cumulative distribution functions (eCDFs) for estimated medians timers and subsequent division times of $CD4^+$ T cells under IL-2 blockade. Each step-up represents an individual donor (HD: green, CeD: red). (B) Permutation test results and null distributions for $CD4^+$ T cells. Histogram show test statistics from $10^7$ random permutations; vertical dashed line indicates the observed statistic, with two-sided p value shown. (C, D) eCDFs (C) and permutation test (D) analyses for CD8+ T cells. (E) Timer medians and subsequent division times were compared between HD and CeD cohorts for $CD4^+$ and $CD8^+$ T-cell responses. Data shown are mean ±SEM of HD and CeD groups ($CD4^+$ N: HD = 22, CeD = 16; $CD8^+$ N: HD = 3,4 CeD = 15). Comparisons used unpaired t-tests with Welch's correction.

#### Supplementary Figure 8

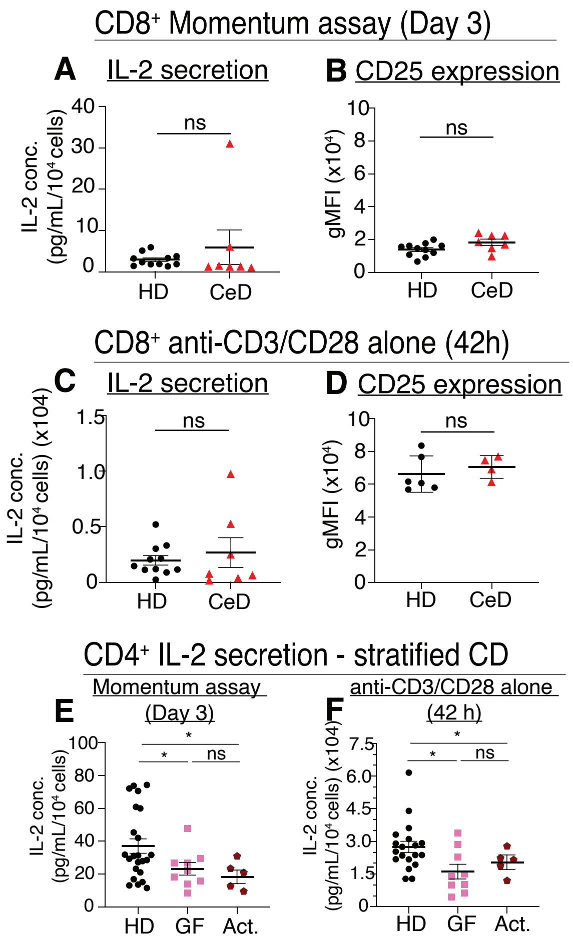


**Supplemental Figure 8. IL-2 secretion and CD25 expression in activated naïve CD8+ T cells and CeD IL-2 subgroup analysis.**

Supernatants and cells were collected from CD4⁺ and CD8⁺ T cell cultures stimulated using the momentum assay at Day 3 or from cultures containing anti-CD3/CD28 beads alone at 42 hours. **(A)** IL-2 concentrations were measured in supernatants and **(B)** cells were stained to measure surface expression of high affinity IL-2Rα (CD25) by flow cytometry at day 3 of the momentum assay (n: HD = 11, CeD = 7). **(C)** IL-2 concentration (N: HD = 11, CeD = 8) and **(D)** CD25 expression (N: HD = 6, CeD = 4) were quantified at 42 h when CD8⁺ T cells where cultured in anti-CD3/CD28 beads alone. **(E)** IL-2 secretion was assessed in cultures containing CD4⁺ T cells from healthy, gluten free- (GF) and active- (Act) CeD at day 3 of the momentum assay (N: HD = 23, GF = 9, Act = 5) and **(F)** 42 h with anti-CD3/CD28 beads alone (N: HD = 19, GF = 9, Act = 5). Data shown as mean ±SEM for Il-2 concentration and CD25 gMFI. Comparisons used unpaired t-tests with Welch's correction. * indicates $p<0.05$

### Supplementary Figure 9

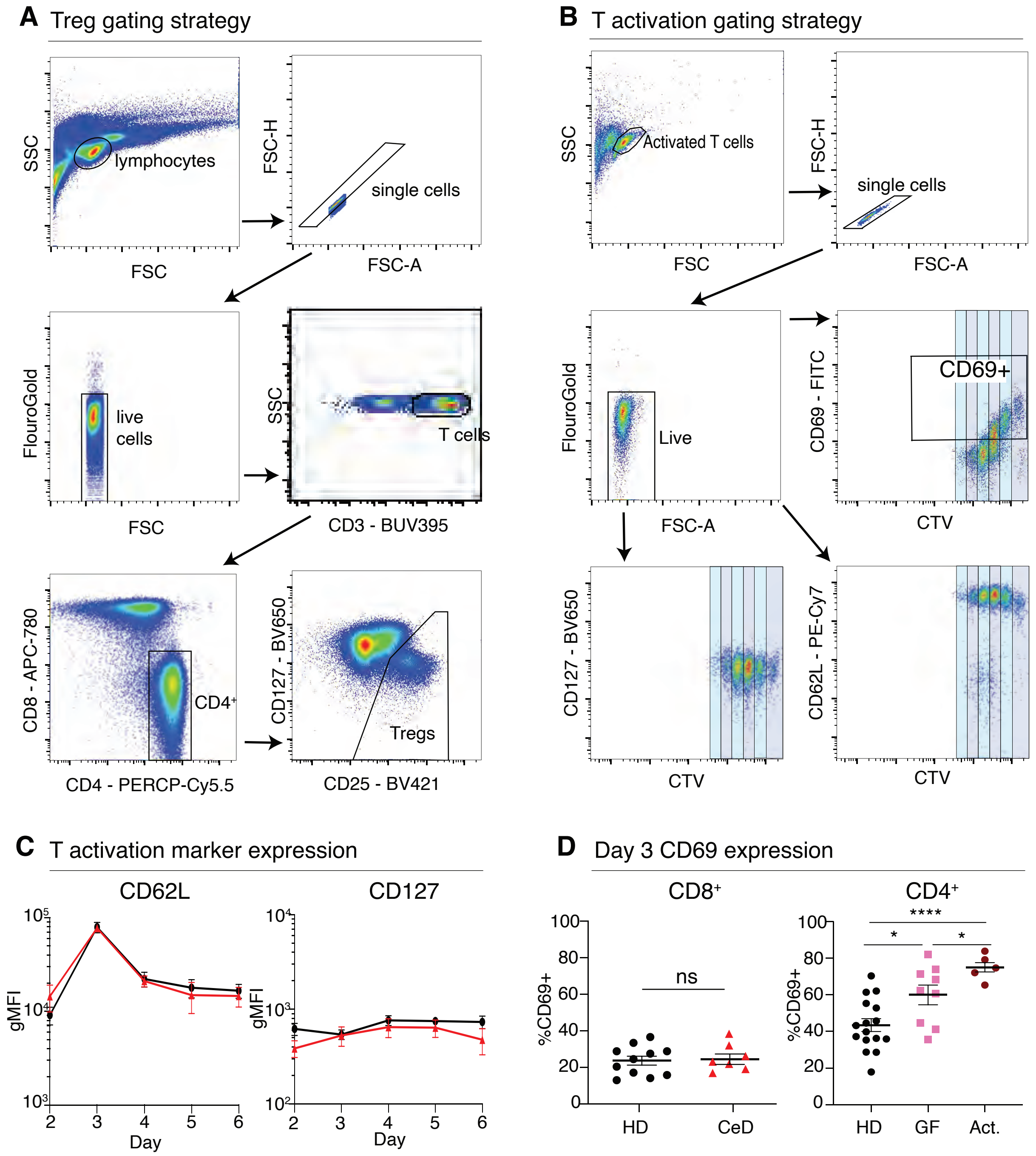


**Supplemental Figure 9. Flow cytometry gating strategies and T cell activation marker analysis**

(A) Treg gating strategy: PBMCs were stained with Fluorogold and antibodies to identify T regulatory cells ($CD3^+CD4^+CD8^-CD25^{hi}CD127^{lo}$). Gating strategy involved gating in the order of lymphocytes, single cells, live cells, $CD3^+$ T cells, $CD4^+CD8^-$ cells, $CD25^{hi}CD127^{lo}$ cells. **(B)** T cell activation marker gating: naïve $CD4^+$ T cells stimulated via the T cell momentum assay were stained for CD69, CD25, CD62L and CD127, followed by live/dead dye (Fluorogold). Gating strategy involved gating in the order of lymphocytes, single cells, live cells, then CTV vs each activation marker (CD69, CD25, CD62L or CD127). **(C)** gMFI of CD62L (left) and CD127 (right) was quantified across the response (N: HD = 8-9, CeD = 9). **(D)** $\%CD69^+$ cells on day 3 of the of the momentum assay for CD8+ T cells (left; N: HD = 11, CeD = 7) and CD4+ T cells (right) was determined for healthy, GF and active CeD individuals (N: HD = 16, GF = 9, Act = 5). Data presented as mean ±SEM. Comparisons used unpaired t tests with Welch's correction. * indicates $p<0.05$; **** indicates $p<0.0001$
